# Orally Administered Exosomes Alleviate Mouse Contact Dermatitis through Delivering miRNA-150 to Antigen-Primed Macrophages Targeted by Exosome-Surface Antibody Light Chains

**DOI:** 10.1101/2020.07.22.214866

**Authors:** Katarzyna Nazimek, Krzysztof Bryniarski, Wlodzimierz Ptak, Tom Groot Kormelink, Philip W. Askenase

## Abstract

We previously discovered suppressor T cell-derived, antigen (Ag)-specific exosomes inhibiting mouse hapten-induced contact sensitivity effector T cells by targeting antigen-presenting cells (APCs). These suppressive exosomes acted Ag-specifically due to a coating of antibody free light chains (FLC) from Ag-activated B1a cells. Current studies aimed at determining if similar immune tolerance could be induced in cutaneous delayed-type hypersensitivity (DTH) to the protein Ag (ovalbumin, OVA). Intravenous administration of a high dose of OVA-coupled, syngeneic erythrocytes induced CD3^+^CD8^+^ suppressor T cells producing suppressive, miRNA-150-carrying exosomes, also coated with B1a cell-derived, OVA-specific FLC. Simultaneously, OVA-immunized B1a cells produced exosome subpopulation, originally coated with Ag-specific FLC, that could be rendered suppressive by in vitro association with miRNA-150. Importantly, miRNA-150-carrying exosomes from both suppressor T cells and B1a cells efficiently induced prolonged DTH suppression after single systemic administration into actively immunized mice, with the strongest effect observed after oral administration. Current studies also showed that OVA-specific FLC on suppressive exosomes bind OVA peptides, suggesting that exosome-coating FLC target APCs by binding to Ag-major histocompatibility complexes. This renders APCs able to inhibit DTH effector T cells. Thus, our studies described a novel immune tolerance mechanism mediated by FLC-coated, Ag-specific, miRNA-150-carrying exosomes that are particularly effective after oral administration.

## 1. Introduction

Recent studies have shown that immune regulation in vivo is more diverse than previously appreciated. A unique antigen (Ag)-specificity of T cell immunosuppression was described previously in contact sensitivity (CS) induced by epicutaneous immunization of mice with reactive hapten Ags [1-5]. This form of Ag-specific T cell tolerance is systemically generated by intravenous (IV) administration of high doses of hapten-conjugated syngeneic erythrocytes, followed by skin sensitization with the same reactive hapten Ag. This induces CD8^+^ suppressor T cells (Ts), that are not FoxP3^+^ regulatory T (Treg) cells [4].

The Ts cells produce and release suppressive, Ag-specific extracellular vesicles (EVs), namely exosome-like nanovesicles [4], that deliver inhibitory miRNA-150. Exosome-targeted antigen-presenting cells (APC), in turn, suppressed CS effector T cells [4,6]. Exosome-mediated suppression was unequivocally confirmed by an in vivo experiment, showing that systemic administration of these exosomes to actively sensitized hosts at the peak of the hapten-specific effector T cell-mediated CS strongly reduced subsequent immune skin swelling responses measured for over four days [4].

The Ag-specificity of this Ts cell-derived exosome-mediated suppression is due to a surface coating with antibody (Ab) free light chains (FLC) provided by B1a cells that also were activated during tolerogenesis [4]. This Ag-specificity of suppressive exosomes was proved in dual Ag crisscross experiments using two non-cross reactive haptens, and also in similar dual crisscross Ag-affinity column chromatography [4].

We showed previously that contact immunization activates peritoneal B1a cells to migrate to the spleen, where they produce specific IgM Ab and their derived Ab FLC [7]. By tolerizing µMT and JH^neg/neg^ antibody deficient mice, and coating the resulting exosomes with chosen FLC, we demonstrated that the Ag-specific Ab FLC bind to the surface of Ts cell-derived exosomes. This in turn is responsible for the Ag-specificity of the exosome targeting of CS effector cells [8]. Further, the B1a cells also release non-suppressive exosomes, that already express Ag-specific B cell receptor (BCR) and/or surface Ab FLC [9]. Consequently, we have uniquely shown that in vitro association of these Ag-specific B1a cell-derived exosome-like nanovesicles with miRNA-150, but not control miRNAs, also renders them suppressive. Their effect is similar to the one induced by the exosomes derived from the Ts-cells of Ag-tolerized mice, that endogenously acquired miRNA-150. This was termed “an alternate Ag-specific exosome-mediated suppression pathway” [10].

Results of these prior studies on CS suppression led to the conclusion that in vivo these Ag-specific B1a cell-derived exosome-like nanovesicles can associate with exogenous inhibitory non-exosomal miRNA, perhaps carried in vivo by RNA-binding argonaute proteins [4]. Consequently, such miRNA-150-associated exosomes were strongly inhibitory against CS effector cells [4]. Therefore, it seemed that this miRNA likely was acquired from the freely circulating, extracellular pool of non-exosomal RNAs protected from RNases by chaperones like Argonaut proteins [4]. As an in vivo consequence, these B1a cell-derived exosomes seem able to act indirectly by similar Ag-specific binding to surface Ag of APCs, that then inhibit CS-effector T cells via transfer of their acquired inhibitory miRNA-150 [10].

Given the demonstrated initial suppressive exosome targeting of the APC in CS [6], it was postulated that the actual Ag bound by these FLC-coated suppressive exosomes could be hapten chemically conjugated to self-protein-peptides complexed with MHC on the APC surface. In the current study, we sought to determine if similar suppression mechanisms applied to classical cutaneous delayed-type hypersensitivity (DTH) induced by ovalbumin (OVA) protein Ag, and clinically referred to as contact dermatitis. In this case, it was postulated that exosome-coating FLC might bind target OVA Ag peptide determinants complexed with MHC on the APC surface. The most definitive experiments were administering suppressive exosomes systemically to actively immunized mice at the peak of the skin responses and determining effects on elicited DTH over subsequent days [4]. Surprisingly, the resulting long term systemic suppression resulting from oral administration of either the T or B cell-derived suppressive exosomes was superior to intravenous (IV) or intraperitoneal (IP) injections.

## 2. Results

### 2.1. Intradermal immunization with OVA induces DTH effector cells

Mice were immunized for induction of protein Ag-specific DTH by ID injections of a high dose of OVA (100 µg) without an adjuvant. This allowed elicitation of classical late 24 hour DTH responses to ID ear injection (challenge) with a low dose of OVA (10 µg) on day four (Protocol, Figure S1A), compared to controls immunized just with PBS and again similarly ear challenged with OVA (Protocol, Figure S1B). The DTH response consisted of an initial, weakly Ag-specific, early edematous ear swelling component, peaking at 2 hours after challenge (Figure S1C), shown previously to be due to B1a cell-derived IgM Ab and their derived Ab FLC [9,11]. This early component is required for elicitation of the subsequent classical, Ag-specific, late inflammatory phase, that peaks 24 hours after skin challenge (Figure S1D) [7,9,11]. Three OVA ID immunizing doses were used (20, 100 and 500 µg). For this dose range, the intensity of elicited DTH, measured as ear swelling response [12], was similar (Figure S1D, Groups B, C and D vs. A). Thus, we chose 100 µg of OVA in saline as the optimal immunizing dose administered by ID injection into 4 sites, as previously described [11].

### 2.2. Adoptively transferred DTH-effector cells were strongly inhibited by incubation in vitro with Ts cells that produced suppressive exosomes

Immune tolerance was induced by IV injection on day 0 and 4 of high doses of OVA Ag linked to syngeneic red blood cells (RBC), followed by OVA ID injection on day 8 and 9 (Protocol, Figure S2A). As tolerogen, OVA was coupled to syngeneic RBC in the presence of either Chromium Cl (Cr^3+^) or EDC (Figure S2B, 53% suppression), and EDC was chosen as the optimal coupling agent.

Inhibition of DTH by the tolerogenic procedure was found to be due to generation of Ts cells. One way to show this was by adoptive transfer of DTH effector cells harvested from the ID optimally immunized mice that often were employed as a positive response for testing of Ts cell suppressive exosome activity (Protocol, Figures S2C and S2D, left half). Their suppressor activity was determined by in vitro combining and briefly incubating the Ts cells, or their derived exosomes, with the DTH-effector cells just before the adoptive transfer. Suppression was assayed by inhibition of the co-adoptively transferred OVA DTH-effector cells. This was quantitated by measurement of elicited ear swelling activity of the DTH-effector cells in naive recipients that were ear challenged with a low dose of OVA (Protocol, Figure S2D, right half).

### 2.3. Ts cell suppression was due to production of suppressive EVs that by surface expression of tetraspanins are common classical exosomes, and uniquely also co-express surface Ab FLC

The Ts cells were collected on day 11 by harvesting and pooling lymph node and spleen cells from the tolerized mice. These cells were then cultured for 48 hours to yield supernatant containing the suppressive exosomes (Figure 1A). This combination of IV and ID Ag tolerizing injections of OVA generated OVA-specific Ts cells that released OVA-specific suppressive exosomes able to inhibit adoptive transfer of the DTH effector cells (Figure 1A, Group B vs. A, 80% suppression). Furthermore, we previously showed in hapten-induced CS that the Ts cells from hapten tolerized mice are CD3^+^ and CD8^+^ [4]. Similarly in the current OVA DTH system, there also was loss of exosome-suppressive activity when either CD3^+^ or CD8^+^ populations from the lymph node and spleen cells of mice tolerized with IV OVA-RBC, were depleted with specific IgG monoclonal antibodies (mAb) and complement (Figure 1B).

**Figure 1.**
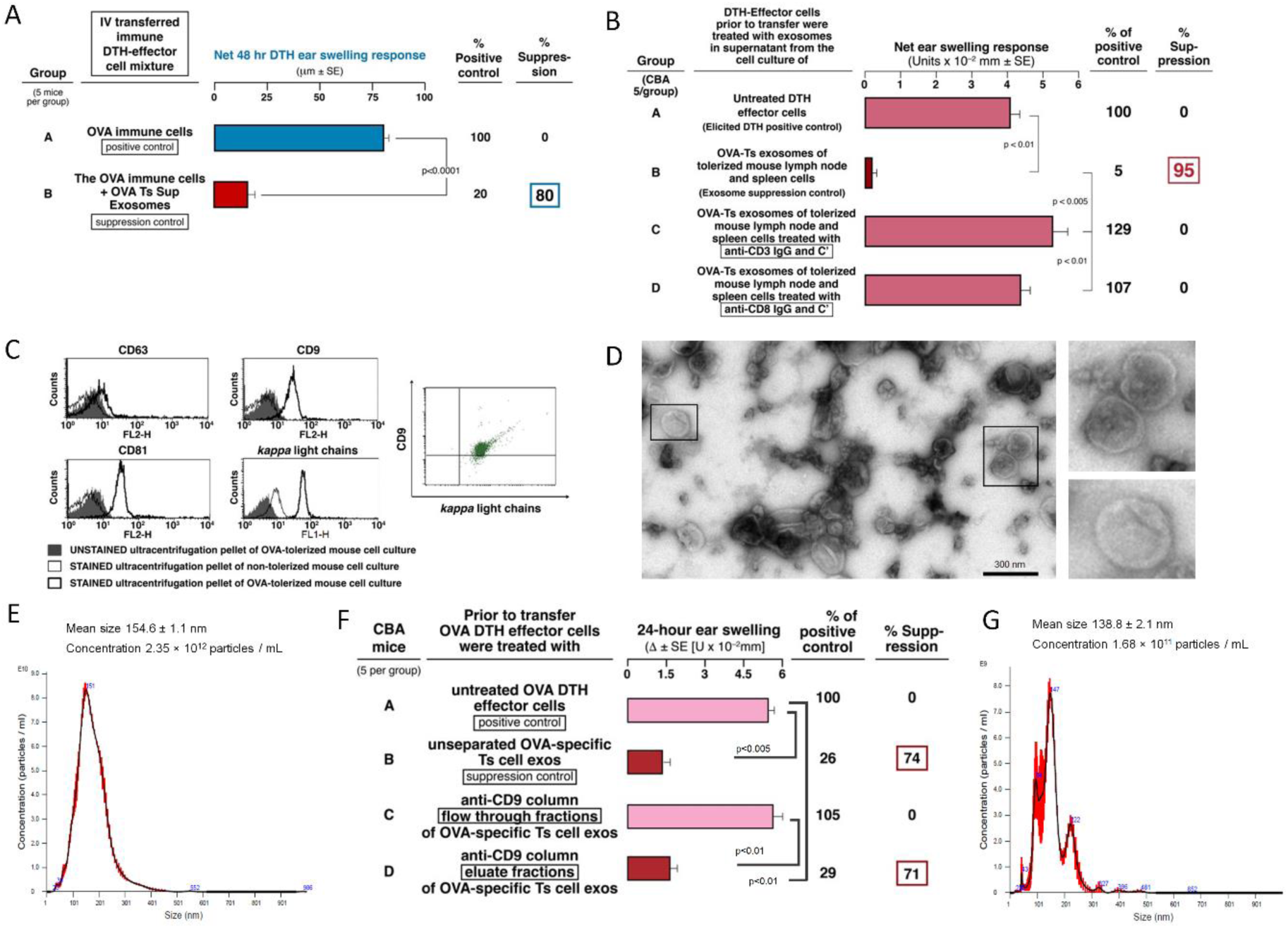
OVA Ag-specific Ts cell-derived, CD9^+^ exosomes inhibit adoptive transfer of T cell mediated DTH. (A) OVA Ag-specific suppressor T cell-derived exosomes inhibit OVA-immune DTH effector cells (Group B, 80% suppression) undergoing adoptive transfer of DTH (Group A, immune response control). (B) Lymph node and spleen cells of Ag-tolerized mice failed to produce suppressive exosomes when depleted of CD3^+^ (Group C) and CD8^+^ (Group D) cell populations vs. their exosome suppression control (Group B, 95% suppression). (C) Flow cytometry showing that OVA-specific Ts cell-derived exosomes express CD9, CD63 (weakest expression) and CD81 tetraspanins, as well as antibody kappa light chains (left and center), and simultaneously co-express CD9 and antibody light chains (right). (D) OVA-specific suppressive exosomes were analyzed with transmission electron microscopy, that showed the presence of nanovesicles with bilamellar membranes. (E) Nanoparticle Tracking Analysis of OVA-specific Ts cell-derived exosomes to estimate their size and concentration. (F) OVA-specific suppressive exosomes were separated onto an anti-CD9-affinity chromatography column. The resulting anti-CD9 binding and non-binding exosome fractions were then used to treat OVA DTH effector cells prior to their adoptive transfer into naive recipients that 24 hours later were challenged with OVA. The elicited DTH ear swelling responses were measured 24 hours later with an engineer’s micrometer, showing that the full suppressive activity of the Ts cell-derived exosomes was in the anti-CD9 binding fraction and there was no suppressive activity in the non-binding flow through fraction (Group D vs. C, 71% suppression vs. 0% suppression). (G) Nanoparticle Tracking Analysis of OVA-specific Ts cell-derived exosomes present in the eluate fraction resulting from separation onto an anti-CD9 affinity chromatography column, to estimate their size and concentration. Results of in vivo assays are shown as delta ± standard error (SE), one-way ANOVA with post hoc RIR Tukey test. *n* = 5 mice in each group or 3 samples in each experimental repetition.

In addition to biologic activity, the descriptive characteristics and properties like phenotype of the assayed OVA Ag-specific suppressive exosome vesicles resembled the hapten-specific exosome-like nanovesicles isolated from mice tolerized to hapten also produced by CD3^+^ and CD8^+^ T cell populations [4]. OVA-specific exosomes highly expressed CD9 and CD81 tetraspanins with significantly lower expression of CD63 (Figure 1C). Noteworthy for the current OVA system, all of the exosome vesicles expressing CD9 tetraspanin also co-expressed surface Ig FLC (Figure 1C, bottom center and right). In addition, transmission electron microscopy revealed the presence of vesicles with billamelar membrane (Figure 1D), and of small size determined by Nanoparticle Tracking Analysis (Figure 1E). Importantly, these Ts cell-derived exosomes were determined to be biologically active by isolation and then testing for suppressive function of the subpopulation eluted from anti-CD9-linked affinity columns that was strongly suppressive compared to the flow through fraction that did not bind to the anti-CD9 columns (Figure 1F, Group D vs. C). Nanoparticle Tracking Analysis of OVA Ts cell exosomes eluted from the anti-CD9 affinity column proved the presence of small vesicles in the column eluate (Figure 1G). Thus, special phenotypic characteristics of small exosome vesicles was surely linked to their function. The linkage of phenotype and function is not often tested elsewhere. In addition, when OVA-specific Ts cell-derived exosomes were separated on OVA-linked affinity column, they also strongly suppressed adoptively transferred DTH response (Figure 2A).

**Figure 2.**
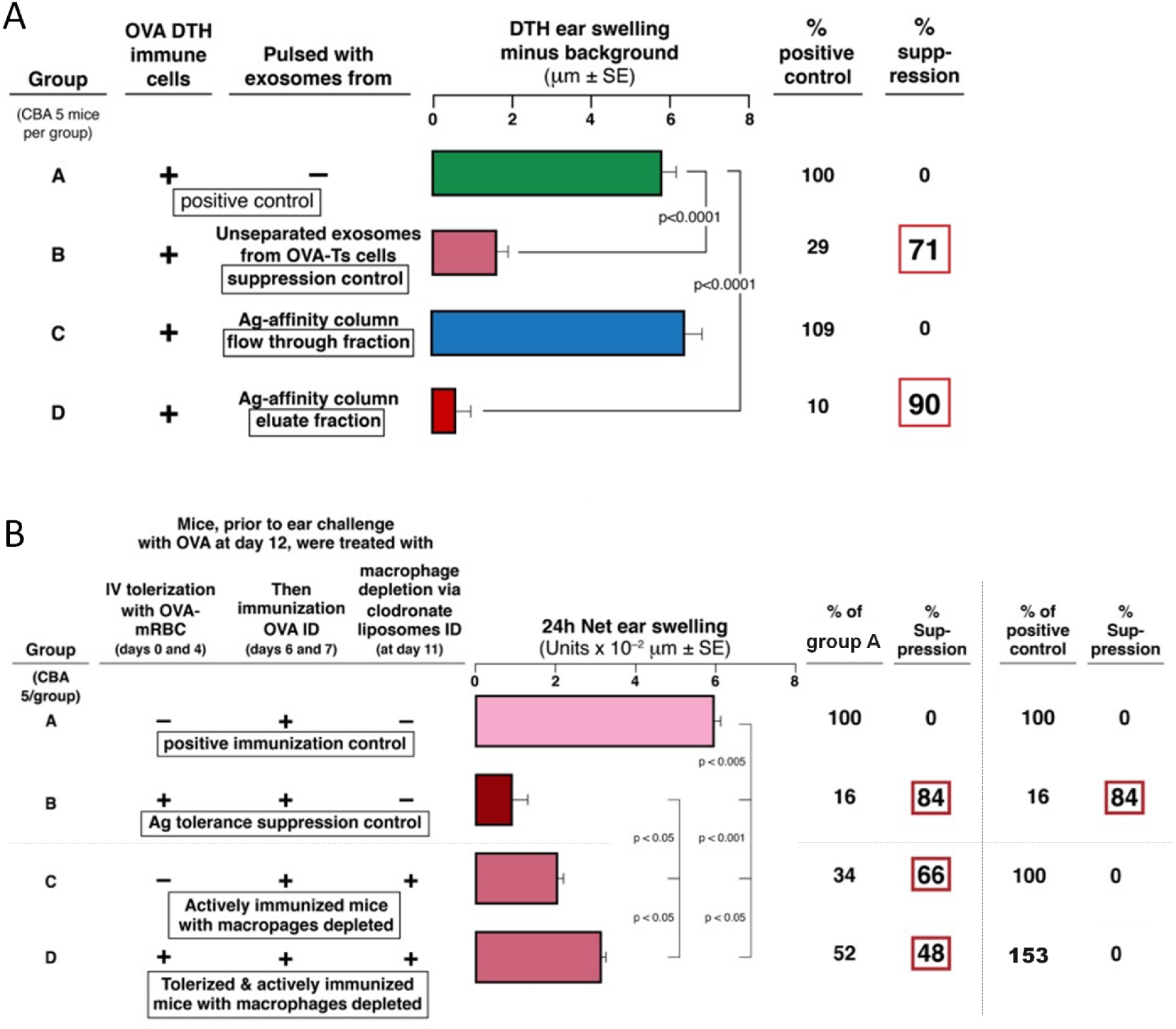
OVA Ag affinity chromatography of the OVA-specific, suppressive exosomes and the essential role of antigen-primed macrophages in tolerance induction. (A) OVA-induced DTH is suppressed only by exosomes present in the eluate of OVA Ag-affinity chromatography column separating the exosomes (Group D column eluate vs. Group C column flow through, 90% suppression vs. 0% suppression). (B) OVA-induced DTH cannot be suppressed by intravenous tolerization with systemic administration of high doses of OVA-linked syngeneic red blood cells in mice depleted of antigen-presenting macrophages (Group D vs. C). Results are shown as delta ± standard error (SE), one-way ANOVA with post hoc RIR Tukey test. *n* = 5 mice in each group.

In summary, all of these findings concerning phenotypic and descriptive properties of the OVA Ts exosomes, fit with the recent most thorough description of true classical exosome characteristics [13]. We also have linked this phenotype to the suppressive function of EVs by testing of their biological activity after separation on OVA Ag and anti-CD9 affinity columns (Figures 1F and 2A). Therefore, we have classified the OVA-specific, Ts cell-derived, functionally active, suppressive nanovesicles as exosomes, a small subtype of the vast family of EVs.

### 2.4. The role of antigen-presenting macrophages in exosome-mediated suppression

Ts cell-derived exosomes that had bound to OVA Ag-linked sepharose columns were found to suppress DTH effector cells (Figure 2A), suggesting that APCs may be targeted by these exosomes. Accordingly, previous studies on suppression of hapten-induced CS showed that Ag-presenting macrophages are targeted by the Ts cell-derived exosomes, that in turn inhibited the DTH-effector T cells [6]. Thus, the Ts cell-derived exosome mediated suppression depended on the presence of macrophages acting as APC for the effector T cells [6]. In the current study, we have similarly observed that depletion of macrophages abolishes the suppression in mice tolerized by IV injection of high doses OVA-RBC (Figure 2B, Group C vs. D). This confirms that the crucial role of Ag-presenting macrophages also applies to the exosome-mediated suppression of OVA-induced DTH.

This experiment shows further that depletion of macrophages also caused decreased DTH-ear swelling responses in actively OVA DTH immunized mice (Figure 2B, Group C vs. A). Therefore, this additionally demonstrates the important role of macrophages acting as APC for the development of the optimal late effector phase of OVA-induced DTH. Thus, for calculations, considering this decreased response in macrophage-depleted actively sensitized hosts as a 100% DTH response, similar macrophage depletion before the elicitation of DTH in the animals that were OVA Ag-tolerized IV prior to the second step of ID immunization, caused a complete reversal of that suppression (Figure 2B, Group D vs. C). Therefore, these data indicate that the tolerogenesis and, as expected, the ability to suppress active DTH, both depend on Ag-presenting macrophages.

### 2.5. OVA-specific Ts cell-derived suppressive exosomes can strongly inhibit active OVA-induced DTH in vivo

Besides demonstrating the suppression of the adoptively transferred DTH effector cells in vitro mixed with Ts cells or their exosomes, we conducted clinically oriented in vivo experiments. For this purpose, we injected the exosomes derived from the OVA tolerized Ts cells directly into mice with active OVA-induced DTH. When systemically administered into mice via the IP route at the time of immunization, the exosomes were only weakly and briefly inhibitory (Figure 3A). In contrast, they had far greater suppressive activity when injected IP into actively immunized mice just before elicitation of DTH on day 4 (Figure 3B). Most important were results obtained when systemic IP exosome treatment began just after determining the peaking 24 hour responses (Protocol, Figure 3C). In kinetic testing, we found that the OVA-specific, Ts cell-derived exosomes strongly suppressed subsequent daily continuing DTH responses at 48, 72, 96 and 120 hours after IP administration of a single physiological dose (Figure 3D). This experiment showed the clinically applicable in vivo suppressive activity of the OVA-specific Ts cell-derived exosomes.

**Figure 3.**
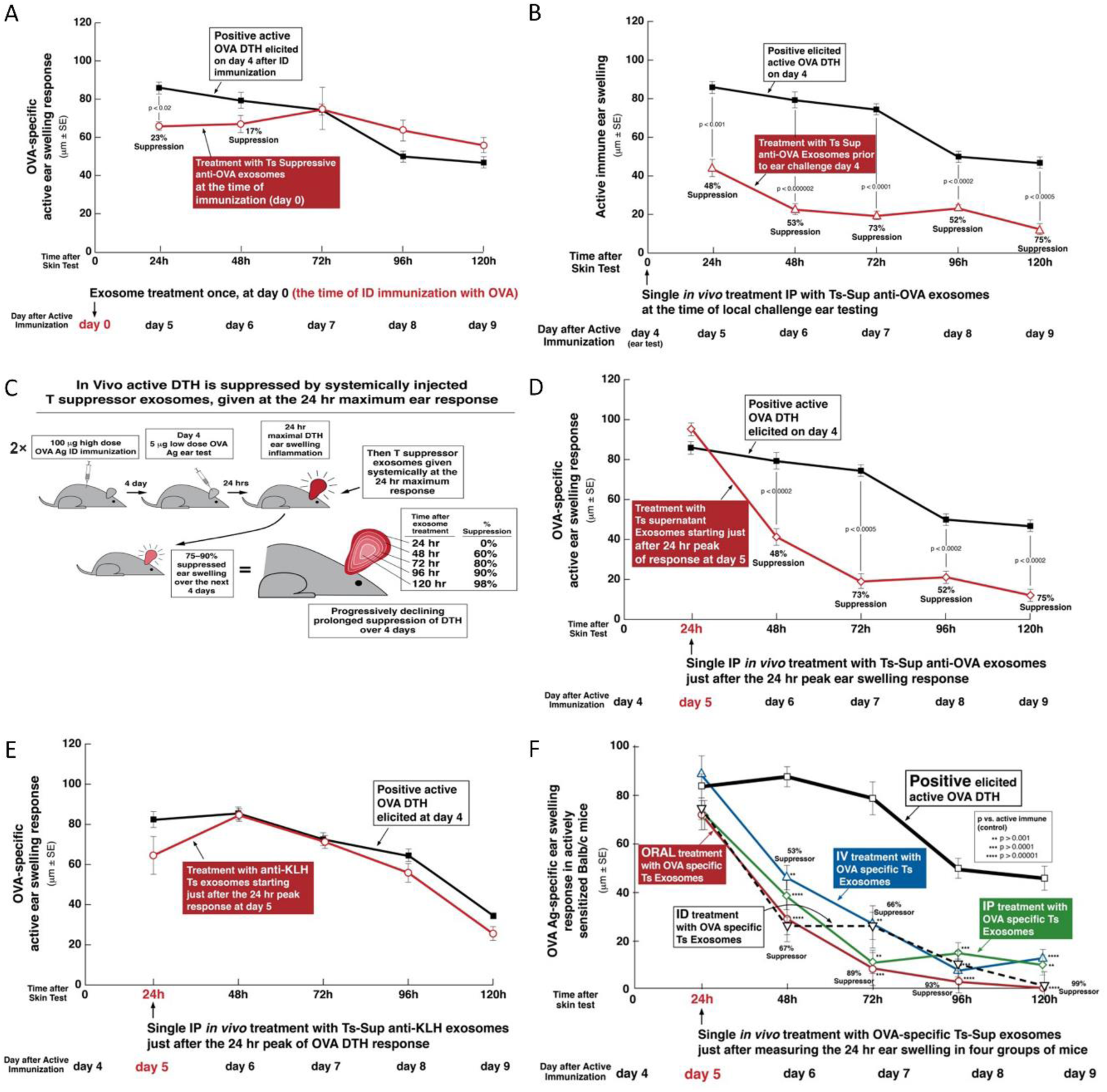
In vivo treatment of active OVA DTH with a systemically administered, single dose of suppressor T cell-derived exosomes, given at various times, and by various routes, including oral and intradermal (ID), with assessment of their Ag-specificity. (A) In vivo treatment of OVA-induced, active DTH with OVA-specific, suppressive exosomes, administered IP at the time of immunization on day 0, that were not suppressive of subsequently elicited 24 hour DTH ear swelling responses measured over five days. (B) In vivo treatment of OVA-induced, active DTH with OVA-specific, suppressive exosomes, administered IP just before elicitation of the DTH response by ear challenge on day 4, that were strongly suppressive (48-75% suppression over the subsequent five days). (C) Protocol: In vivo treatment of active OVA DTH with systemically administered suppressor T cell exosomes, given just after measurement of the 24 hour maximum ear swelling response. (D) In vivo intraperitoneal (IP) treatment of OVA active DTH with OVA-specific suppressor T cell-derived exosomes administered at the 24 hour peak of response, that were strongly suppressive (48-75% suppression over the subsequent four days). (E) Ag specificity of suppressive exosome treatment employing analogous anti-KLH protein suppressor T cell exosomes from KLH Ag tolerized donors, that do not inhibit OVA DTH when given IP at the 24 hour peak of the OVA-elicited DTH ear swelling response. (F) Comparing intravenous (IV), IP, ID and oral treatment of actively OVA DTH immunized mice with OVA-specific suppressor T cell-derived exosomes; starting at the 24 hour peak of response, and measured over the subsequent four days; all significantly inhibited DTH ear swelling; especially oral treatment (the red line). Results are shown as delta ± standard error (SE), one-way ANOVA with post hoc RIR Tukey test. *n* = 5 mice in each group.

### 2.6. In vivo testing of Ag-specificity of the Ts cell-derived exosomes

Similar Ts cell-derived exosomes, from a comparable control DTH system induced by the antigenically non-cross reactive protein keyhole limpet hemocyanin (KLH) showed that the OVA exosome suppression was Ag-specific. In this case, KLH-specific exosomes harvested from lymphoid cells of mice identically tolerized to KLH, and similarly IP injected into actively immunized mice just after the peak of the active 24 hour OVA-induced DTH response, were totally inactive (Figure 3E). In contrast, Ts cell-derived exosomes harvested from mice tolerized by IV injection of high doses of OVA-RBC (Figure S2A), Ag-specifically inhibited DTH in mice actively immunized with OVA, when delivered systemically IP just before (Figure 3B), and, most importantly, 24 hours after ear challenge with Ag (Figure 3D). It thus seems that Ag-specificity of exosome mediated suppression likely resulted from the presence of antibody kappa light chains on the Ts-derived exosome surface that enabled Ag-specific binding to the OVA Ag-linked sepharose in the affinity column (Figures 1C and 2A) [8].

### 2.7. Comparison of IV, IP, ID and PO systemic treatments with OVA Ts cell-derived exosomes, beginning just after the 24 hour peak of OVA DTH, in actively immunized mice

In the studies described above, OVA Ag-specific suppressive exosomes administered IP to actively immunized mice significantly suppressed the in vivo manifestations of OVA DTH reactions (Figures 3B and 3D). This systemic administration of the suppressive exosomes at a single dose by the IP route, had not been compared to any other means of treatment. Therefore, we performed new experiments to test the efficacy of treatment with Ts cell-derived, OVA Ag-specific, suppressive exosomes given by more clinically applicable routes; compared to IP treatment. Thus, these OVA-specific suppressive exosomes again were administered systemically at a single dose judged to be physiological, via different routes at the 24 hour peak of response in mice that were actively OVA immunized to elicit DTH.

Administration of these exosomes via IP route again was quite suppressive over the subsequent 4 days (Figure 3F, green line) compared to positive control, i.e. untreated active DTH (Figure 3F, thick black line). Further, administering the exosomes IV also caused prolonged inhibition (Figure 3F, blue line), but overall slightly less effective compared with the IP route.

Outstandingly, the most efficient suppression was observed after PO administration of the single dose of the inhibitory exosomes (Figure 3F, red line). Finally and interestingly, when treatment with the suppressive exosomes was by the ID route in the torso, this also caused suppression of the DTH responses (Figure 3F, dashed black line). Note that this was via the same route used for Ag immunization, but at different skin sites that also were distant from DTH elicited in the ears.

### 2.8. Non-suppressive Ts cell-derived exosomes from tolerized miRNA-150^neg/neg^ mice are rendered suppressive by association with miRNA-150

In the similar, previously studied tolerance system of the hapten-specific CS response, it was shown that miRNA-150 was required to be carried into the targeted cells by the suppressive exosomes for inhibitory activity [4]. We tested this point here by using exosomes generated by OVA-Ag-tolerized miRNA-150^neg/neg^ mice. Suppression of DTH did not occur in recipients of wild type (WT) OVA-immune DTH-effector cells pretreated in vitro with Ts cell exosomes from tolerized miRNA-150^neg/neg^ mice (Figure 4A, Group C vs. A), compared to the strong inhibition by suppressive exosomes generated from miRNA-150^pos/pos^ wild type donors (Figure 4A, Group B vs. A). This indicated that as in the hapten Ag CS system, DTH suppression caused by tolerization with high doses of Ag generated Ts cell exosomes that were also dependent on their ability to deliver miRNA-150 to targeted cells.

**Figure 4.**
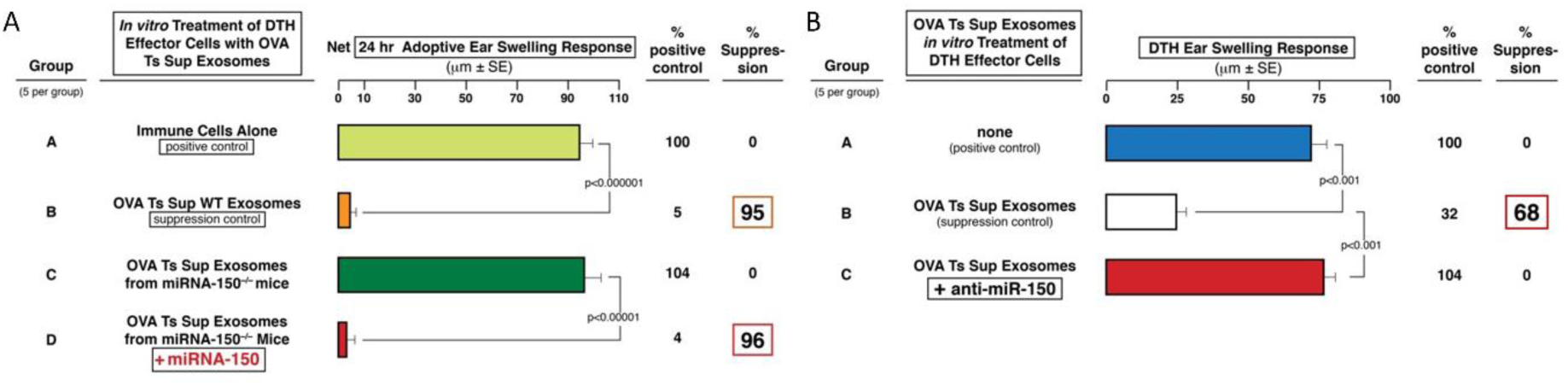
DTH suppression by exosomes from OVA Ag-tolerized mice depends on miRNA-150. (A) Non-suppressive exosomes from OVA Ag-tolerized miRNA-150^neg/neg^ mice were restored for ability to suppress 24 hour OVA DTH by preliminary in vitro association with miRNA-150 (Group D vs. C). (B) OVA Ag-specific suppressor T cell-derived exosomes that suppress adoptive transfer of OVA DTH, are inhibited by pre-incubation with anti-sense polynucleotide to miRNA-150 (anti-miR-150, Group B vs. C). Results are shown as delta ± standard error (SE), one-way ANOVA with post hoc RIR Tukey test. *n* = 5 mice in each group.

In the CS system, the suppressive exosome from hapten Ag tolerized mice, additionally were activated to be able to associate with a given miRNA in vitro. Accordingly, we tested these non-suppressive exosomes from tolerized miRNA-150^neg/neg^ mice after incubation in vitro with synthetic miRNA-150. These miRNA-150-treated exosomes from tolerized miRNA-150^neg/neg^ mice were converted to be able to suppress the adoptively transferred DTH effector cells, like the exosomes from Ag-tolerized WT miRNA-150^pos/pos^ mice (Figure 4A, Group D vs. A and C). Thus, the added miRNA-150 seemed to have become associated with the initially non-inhibitory exosomes from miRNA-150^neg/neg^ tolerized donors to render them capable of fully suppressing adoptive transfer of OVA-specific DTH-effector cells, to the same extent as suppressive exosomes from Ag-tolerized WT miRNA-150^pos/pos^ mice (Figure 4A, Group D vs. B).

As a third experimental confirmation of these findings, OVA Ts cell-derived exosomes from WT mice, presumed to contain inhibitory miRNA-150, were pre-incubated with anti-miR-150 polynucleotide (i.e. a chain with the reverse sequence as miRNA-150), that might hybridize with active miRNA-150. Indeed, this led to strong reversal of the suppression mediated by these anti-miRNA-150 pulsed OVA-induced Ts cell exosomes (Figure 4B, Group C vs. B).

Thus, considering the experiments restoring suppression in exosomes from Ag tolerized miRNA-150^neg/neg^ mice, there were two confirmatory results; (i): showing that initially non-suppressive exosomes from tolerized miRNA-150^neg/neg^ mice were reconstituted by association with miRNA-150 (Figure 4A), and (ii): showing that DTH-inhibition by natively suppressive Ts cell exosomes from Ag high dose tolerized WT mice was abolished by association of these competent exosomes with anti-miRNA-150 polynucleotide (Figure 4B). Therefore, these results together constituted strong evidence that OVA Ts cell exosomes derived from OVA Ag tolerized mice mediated suppression of DTH due to their content and likely delivery of inhibitory miRNA-150.

### 2.9. OVA Ag-specific B1a cell-derived exosomes from Ag-immunized mice, that are non-suppressive, can be rendered suppressive by association with miRNA-150

We hypothesized that exosomes need two properties to be suppressive: the Ag specificity due to surface FLC and delivery of miRNA-150. Accordingly, we tested the ability of non-suppressive, Ag-specific B1a cell-derived exosomes from optimally immunized mice, that were non-suppressive, likely because they contained no miRNA-150, to be rendered suppressive by similar association with miRNA-150 (Protocol, Figure 5A). For this experiment, B1a cell-derived exosomes were harvested from culture supernatants derived from lymph node and spleen cells of mice simply immunized ID with OVA. They were harvested at only 2 days after ID immunization when B1a cells produce Ag-specific IgM antibodies and Ab FLC [9,11]. The B1a cell exosomes in this supernatant were hypothesized to likely be OVA Ag-specific due to membrane anti-OVA BCR, and/or a surface coating with anti-OVA Ab FLC, as shown in Figure 5B [8]. We determined that these immune B1a cell-derived exosomes expressed CD9 and CD81 tetraspanins (Figure 5B, left). Additionally, CD9^+^ vesicles were found to co-express mouse kappa FLC, in contrast to samples from non-immunized animals (Figure 5B, right top vs. bottom). Transmission electron microscopy revealed the presence of bilamellar membranous vesicles in the pellet of ultracentrifuged supernatant of OVA-specific B1 cells (Figure 5C).

**Figure 5.**
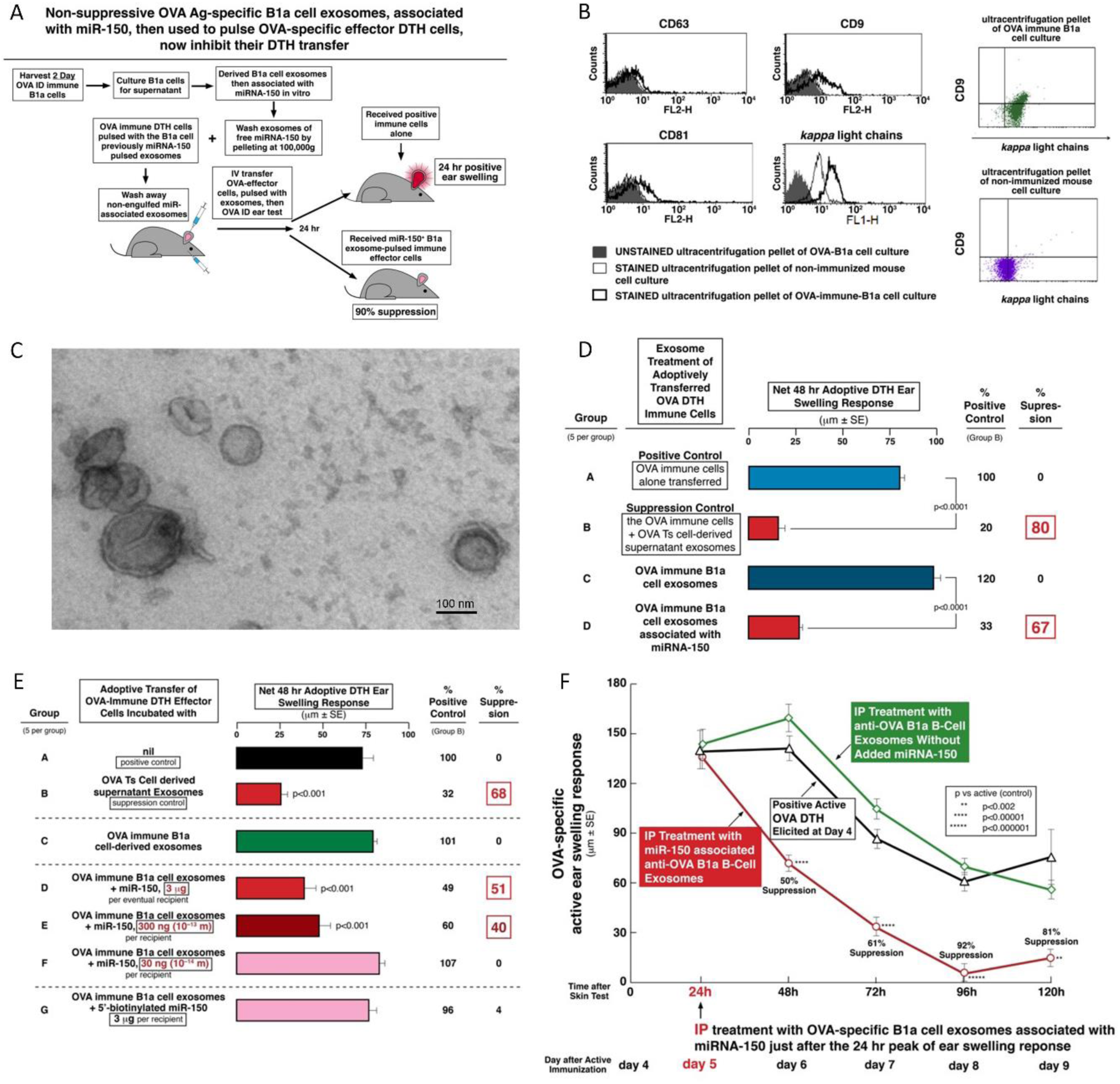
Non-suppressive OVA Ag-specific B1a cell exosomes inhibit adoptively transferred OVA DTH when associated with miRNA-150. (A) Experimental protocol. (B) OVA-specific B1a cell-derived exosomes were cytometrically analyzed for expression of CD9, CD63 and CD81 as general markers of small exosomes, after coupling onto beads (left and center). Further, they were analyzed and found positive for co-expression of CD9 and antibody kappa free light chains (FLC, center and upper right), while some in the pellet from ultracentrifuged culture supernatant from normal lymphoid cells expressed Ab FLC, but did not co-express CD9 (right lower). (C) B1a cell-derived exosomes were analyzed with transmission electron microscope, showing the presence of bilamellar vesicles. (D) Non-suppressive OVA Ag-specific B1a cell-derived exosomes become suppressive of OVA DTH adoptive transfer when in vitro associated with miRNA-150 (Group C vs. D). (E) A dose response experiment shows that non-suppressive OVA Ag-specific B1a cell-derived exosomes are rendered suppressive of DTH effector cell adoptive transfer by their association in vitro with low doses of miRNA-150. (F) OVA Ag-specific B1a cell-derived exosomes associated in vitro with inhibitory miRNA-150 are able to suppress active OVA DTH when injected intraperitoneally (IP) at the 24 hour peak of ear swelling response, in contrast to similar exosomes not associated with miRNA-150 (red line vs. green line, 50-92% suppression over the subsequent four days), and compared to no treatment of this active DTH (the thick black line). Results of in vivo assays are shown as delta ± standard error (SE), one-way ANOVA with post hoc RIR Tukey test. *n* = 5 mice in each group or 3 samples.

As expected, these exosomes derived from optimally immunized, but non-tolerized donors were non-suppressive, likely because they did not naturally contain endogenously generated miRNA-150 because they were not from tolerized mice (Figure 5D, Group C vs. B). In contrast, when these B1a cell-derived, Ag-specific exosomes were in vitro associated with miRNA-150, they became suppressive (Figure 5D, Group D vs. C).

In summary, these data of the OVA protein DTH system confirmed that Ag-specific, suppressive exosomes could be actually derived either by CD8^+^ Ts cells, then they natively contain miRNA-150 and express surface Ag-specific FLC donated by neighboring activated B1a cells (Figure 1A, Group B), or, alternatively, by immune B1a cells, then they natively express Ag-specific FLC and could be further associated with miRNA-150 (Figure 5D, Group D).

### 2.10. Dose response of miRNA-150 association with immune B1a cell-derived exosomes

This type of miRNA association was new, and thus required exploration of physiological relevance by dose response experimentation. In the CS system, this was shown to be mediated by unusually effective, very low doses of B1a cell exosome-associating miRNA-150 [10]. Accordingly, we determined the least dose of miRNA-150 required for switching OVA-specific B1a cell exosomes to suppressive function in vivo. Use of decreasing serial dilutions of miRNA-150 added to exosomes from B1a cells from early two day OVA immunized mice showed that as little as 300ng or 10^−13^ moles, was the limiting dose of miRNA-150 in this system. This amount was sufficient to associate with B1a cell exosomes to mediate highly significant suppression of adoptively transferred OVA DTH mediating immune cells (Figure 5E, Group E vs. C).

An additional feature of this experiment was an attempt to employ chemically 5’-biotinylated miRNA-150 for eventual testing of function of this labeled polynucleotide in suppressing the adoptive transfer of DTH. Such a construct would be useful for detection, quantitation, and kinetic studies of the miRNA-150 carrying suppressive B1a cell-derived exosomes. However, attempted association of the immune activated B1a cell-derived, OVA Ag-specific exosomes with biotinylated miRNA-150, even at the highest dose of 3 µg, was not suppressive of OVA-specific effector cells in adoptive transfer of DTH (Figure 5E, Group G vs. C). This suggested that the native miRNA-150 (or perhaps just its native 5’ end sequence) was required for association with exosomes and/or eventual inhibition of the mRNA targeted by miRNA-150. Therefore, this group served as an excellent miRNA control.

Overall, these studies confirmed the ability of miRNA-150 to associate with OVA Ag-specific immune B1a cell-derived exosomes, as we found previously with these type of exosomes also from early immune, but hapten-Ag-specific B1a cells [10]. The limiting dose here was much greater than in the hapten system, and biotinylating the 5’ end of miRNA-150 prevented association or function in targeted cells.

### 2.11. Prolonged in vivo suppression by systemic treatment of active OVA DTH responses with B1a cell-derived exosomes associated with miRNA-150

Important in any therapy, is attaining a significant duration of the effect; especially in clinical translation to patients. We showed above strong and prolonged in vivo suppression of existing OVA DTH over four days in actively immunized mice treated with a single dose of Ts cell-derived exosomes administered by multiple routes (Figure 3F). Accordingly, we performed a further kinetic study in the OVA DTH system, but now employing the newly described B1a cell-derived exosomes, rendered suppressive by in vitro association with inhibitory miRNA-150 (Figures 5D and 5E). We found that systemic, single physiological-dose IP treatment begun at the 24 hour peak of response with miRNA-150 associated OVA Ag-specific B1a cell exosomes had similar profound and prolonged OVA DTH suppressive effects (Figure 5F, red line, 50-92% suppression), when compared to identical but miRNA-150 unassociated B1a cell-derived exosomes (Figure 5F, green line) and to hosts with elicited active DTH that had no treatment (Figure 5F, black line). Therefore, OVA DTH suppression like that mediated by T cell-derived exosomes from Ag tolerized mice could be constructed from these B cell-derived exosomes.

### 2.12. Comparing the binding of four different anti-OVA mAb and their derived light and heavy chains, as well as polyclonal anti-OVA FLC to various OVA preparations

An important question about DTH inhibition by Ag-specific, suppressive exosomes is the mechanism of suppression; including the mechanism of the Ag-specificity due to a coating with Ab FLC [8]. To examine this we employed a set of four different anti-OVA IgG monoclonal antibodies, where each was rendered into its Ab heavy and Ab FLC. We compared the ability of these constituent chains to whole IgG in dose response binding to native OVA Ag, adhering to wells of enzyme-linked immunosorbent assay (ELISA) plates and serially diluted from 2.5 to 0.078 µg per ml (Figure 6A).

**Figure 6.**
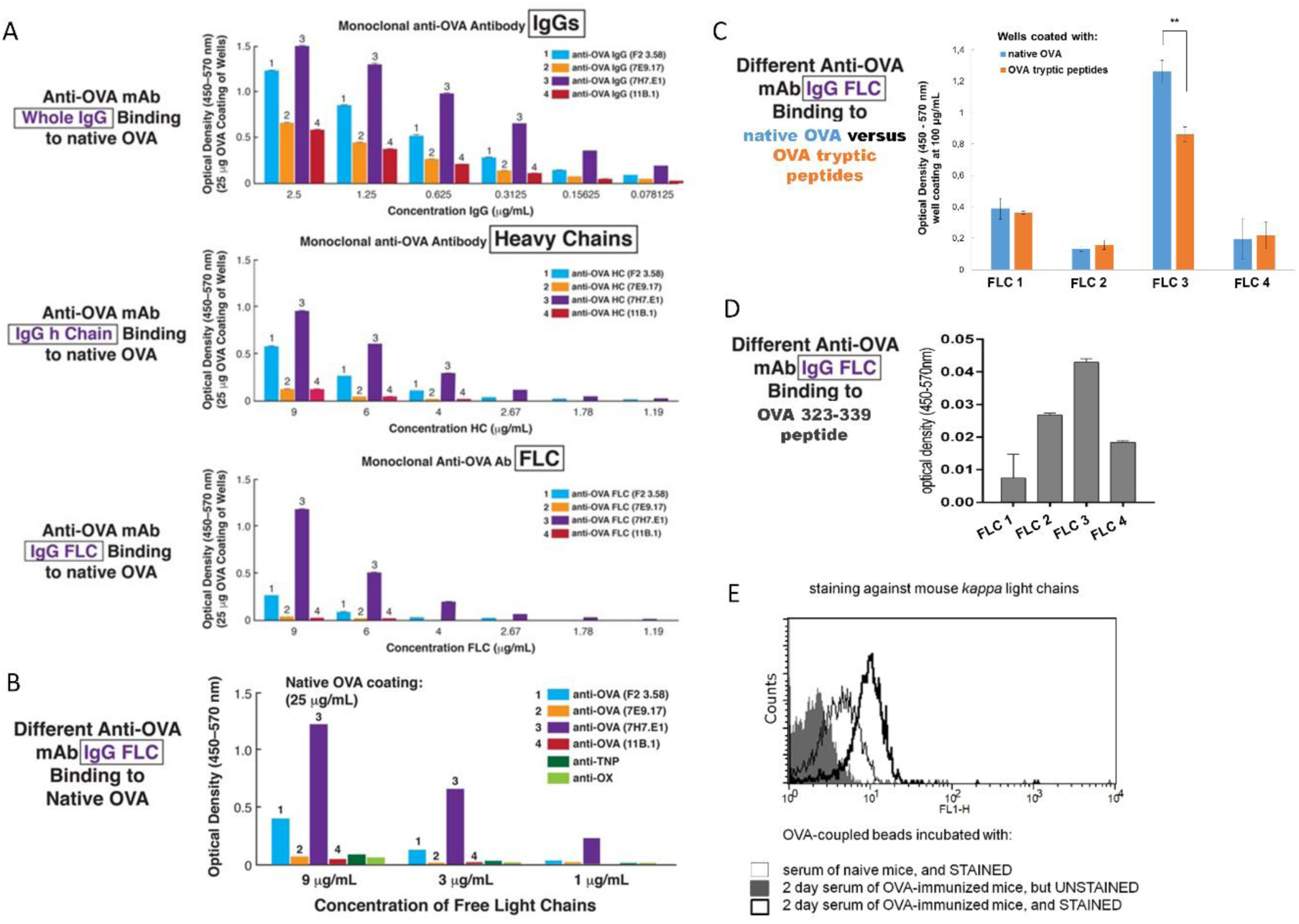
ELISA assays to assess the ability of four different monoclonal anti-OVA Ab-derived free light chains (FLC) to bind to native OVA, OVA tryptic peptides and OVA 323-339 peptide, as well as flow cytometry analysis of anti-OVA Ab FLC in 2-day immune serum. (A) Binding to native OVA by serial dilutions of four different monoclonal anti-OVA Ab-derived FLC and Ab heavy chains, vs. the whole IgGs, shows significant binding with a particular mAb and similar strength with a given mAb and its derived heavy and light chains. (B) Binding of various monoclonal anti-OVA Ab FLC to native OVA compared to binding of control anti-OX and anti-TNP hapten-specific mAb shows the Ag specificity of the Ab FLC binding. (C) Comparison of binding of four monoclonal anti-OVA Ab-derived FLC to either native OVA or OVA tryptic peptides, showing the difference only in the case of FLC #3. The two-tailed Student t test, ** *p* < 0.01. (D) Binding of four monoclonal anti-OVA Ab-derived FLC to OVA 323-339 antigenic determinant. (E) The presence of OVA Ag-specific Ab FLC in 2 day serum of OVA-immunized mice (thick black line) was confirmed by flow cytometry, as compared to serum of control non-immunized mice (thin black line). *n* = 3 wells or samples in each experimental repetition.

Over the dilution range employed, binding to native OVA adsorbed to plastic ELISA wells, occurred with the whole IgGs as well as the separate Ab heavy and light chains. The anti-OVA mAb were numbered #1, 2, 3 and 4, and a rank order of binding was determined. Binding of the isolated heavy and light chains to native OVA Ag was in the same rank order for a given anti-OVA whole monoclonal IgG, and differed in strength of OVA-binding comparing the four mAb (Figure 6A). Binding was best with whole IgG and next the heavy chains and least the FLC (Figure 6A).

Another related ELISA experiment showed that the binding to native OVA by Ab FLC could be claimed Ag-specific, when compared to two different anti-hapten mAb FLC; i.e. anti-oxazolone and anti-TNP FLC (Figure 6B). These various anti-hapten mAb FLC were shown previously to bind to and convert non-suppressive exosomes derived by Ts cells from hapten Ag tolerized pan Ab deficient JH^neg/neg^ mice to suppressive function, in adoptive transfer of a CS-effector cell mixture [4].

In another ELISA assay we have compared the four mAb IgG-derived FLC for their binding to native OVA and OVA tryptic peptides. FLC #1, 2 and 4 bound with a similar strength to both OVA preparations, while binding of FLC #3 to OVA tryptic peptides was weaker than to native OVA (Figure 6C). Furthermore, all assayed FLC were shown to bind to OVA 323-339 antigenic determinant but, however, with a very low efficacy (Figure 6D).

As mention above, antigen-activated B1a cells provide FLC for coating of Ts cell-derived exosomes (Figure 1C). Thus, we speculated that these B1a cell-derived FLC can be found in serum collected from mice immunized with OVA 2 days earlier. Prior studies in the CS system and in a model of early resistance to pneumococcal pneumonia indicated that such sera contain polyclonal anti-hapten IgM Ab, and also Ab FLC with diverse Ag-specificities due to germ line V-region mutated DNA sequences in a subpopulation of these ordinarily germ line expressing early B1a cells [14-16]. Thus, to evaluate this assumption, latex beads coated with native OVA antigen were incubated with OVA immune sera, washed, stained against mouse kappa light chains, and subjected to flow cytometric analysis, that revealed the presence of Ab FLC, which bound to OVA-coated beads, in tested sera (Figure 6E). As expected, the amount of detected anti-OVA FLC was much greater in sera of OVA-immunized mice, when compared to naive mouse sera (Figure 6E).

### 2.13. Testing the anti-OVA mAb FLC that coat Ts cell-derived exosomes reveals their ability to bind to OVA antigenic determinant

To in vivo assess the involvement of Ab FLC in suppression of DTH by Ts cell-derived exosomes, the same amounts of anti-OVA FLC #1, 2, 3 and 4 were used to coat identical aliquots of Ts cell-derived exosomes from OVA Ag tolerized JH^neg/neg^ mice (Figure 7A, protocol). Because of their genetic defect in Ab production, the JH^neg/neg^ mice do not produce native Ab FLC and, therefore, their derived exosomes are not suppressive on their own [4,8,10]. Thus, we tested the relative ability of the four mAb-derived FLC to restore suppressive action of exosomes, when used to coat exosomes derived by Ts cells from OVA Ag-tolerized JH^neg/neg^ mice. We found that the JH^neg/neg^ mouse Ts cell exosomes coated with FLC # 2, 3 and 4 mediated significant suppression of adoptively transferred DTH effector cells (Figure 7B). Simultaneously, we performed an ELISA-based assay to compare the binding of JH^neg/neg^ mouse Ts cell-derived exosomes coated with four mAb-derived FLC to native OVA and OVA tryptic peptides (Protocol, Figure 7C, left), which was found similar in the case of FLC #1, 2 and 4 when tested as freely dispersed in assay buffer (Figure 6C). This assay showed the significantly greater amount of JH^neg/neg^ mouse Ts cell exosomes that bound to OVA tryptic peptides in comparison with binding to native OVA, when these exosomes were coated with FLC #1, 2 and 4, but not with FLC #3 (Figure 7C), suggesting that some FLC bind OVA tryptic peptides stronger when coated onto exosomes. Thus, these observations led us to hypothesize that the ability of Ab light chains to allow exosomes to Ag-specifically suppress DTH effector cells depends mostly on the total avidity of Ag-binding by all FLC that coat suppressive exosomes, which summarizes their individual affinities. To test this hypothesis, we have applied FLC #2 and 3, which differed in their binding capability (Figures 6C and 7C) and suppressive activity (Figure 7B), as well as 2-day serum from OVA-immunized mice, onto affinity chromatography column filled with sepharose conjugated with OVA tryptic peptides. The yielded column eluates containing FLC that had strongly bound to OVA tryptic peptides, were then used to coat JH^neg/neg^ mouse Ts cell-derived exosomes that have been incubated with DTH effector cells prior to their adoptive transfer. We observed that coating of JH^neg/neg^ mouse Ts cell exosomes with OVA tryptic peptide-binding FLC #2 and 3, and OVA-immune serum FLC, led to significant suppression of DTH response (Figure 7D). This confirmed that the ability of FLC to bind the Ag indeed determines its suppressive activity when coating the Ts cell exosomes.

**Figure 7.**
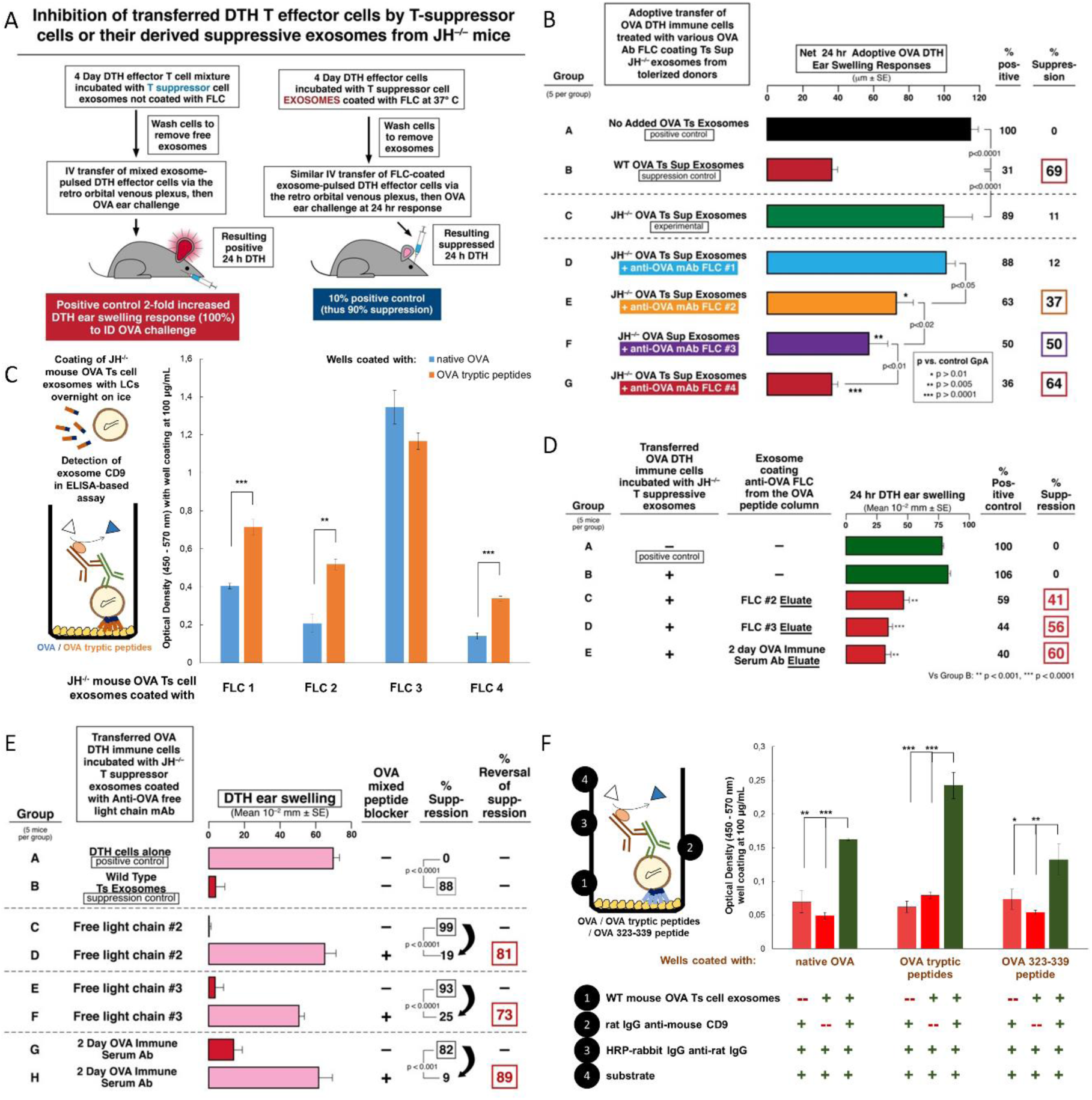
Inhibition of transferred OVA DTH effector cells that were pulsed with Ag-tolerized JH^neg/neg^ mouse exosomes that were in vitro coated with the four different anti-OVA Ab FLC. (A) Protocol: Inhibition of adoptively transferred OVA DTH effector cells when preliminarily pulsed with exosomes from Ag-tolerized WT donors, compared to those from similarly Ag tolerized JH^neg/neg^ mice, that in some cases were additionally coated in vitro with the four various anti-OVA mAb FLC. (B) Adoptive transfer of OVA DTH effector cells pulsed in vitro with OVA-specific Ts cell-derived exosomes from Ag-tolerized JH^neg/neg^ mice, reconstituted for suppression by in vitro coating with the four different anti-OVA mAb FLC. (C) ELISA-based assay (scheme - left) to evaluate the binding of JH^neg/neg^ mouse Ts cell-derived exosomes coated with four different monoclonal anti-OVA Ab FLC to either native OVA or OVA tryptic peptides, showing the stronger binding to OVA tryptic peptides in the case of FLC #1, #2 and #4. (D) Reconstitution of suppression by coating of non-suppressive T cell exosomes derived from OVA Ag-tolerized JH^neg/neg^ mice with anti-OVA mAb FLC #2 and #3 or 2-day immune serum FLC eluted from OVA tryptic peptide-linked affinity column is successful. (E) Use of OVA tryptic peptides to potentially block anti-OVA mAb FLC that coat Ts cell-derived suppressive exosomes. The OVA peptides block anti-OVA mAb FLC and FLC from OVA-immune serum that coated Ts cell-derived suppressive exosomes from Ag-tolerized JH^neg/neg^ mice, on adoptive transfer of DTH, mediated by a mixture of APCs and OVA-specific DTH effector T cells. (F) ELISA-based assay (scheme - left) to evaluate the binding of wild type mouse Ts cell-derived suppressive exosomes to either native OVA, OVA tryptic peptides or OVA 323-339 antigenic determinant, showing that exosomes originally coated with anti-OVA FLC can bind to all assayed OVA preparations. Results of in vivo assays are shown as delta ± standard error (SE, *n* = 5 mice in each group), and of in vitro assays as mean ± standard deviation (SD, *n* = 3 wells in each experimental repetition), one-way ANOVA with post hoc RIR Tukey test or two-tailed Student t test, * *p* < 0.05; ** *p* < 0.01; *** *p* < 0.001.

It has been previously shown that coating of exosomes from Ts cells generated by Ag induced tolerance with Ab FLC mediated the Ag-specificity of suppression [4,6,8]. According to our overall observations, we therefore proposed that FLC, when coating the suppressive exosomes, bind to antigenic determinants, which allows to antigen-specifically target APCs present among DTH effector cells. To initially test this hypothesis, we coated JH^neg/neg^ mouse Ts cell-derived exosomes with FLC #2 and 3, and with serum-derived FLC, and then incubated these FLC-coated exosomes with OVA tryptic peptides prior to treating DTH effector cells. Pre-incubation of FLC-coated exosomes with OVA tryptic peptides blocked their suppressive activity (Figure 7E, groups D vs. C, F vs. E and H vs. G), thereby proving that the interaction of FLC-coated exosomes with an Ag is required for suppression.

Thus, to finally confirm the ability of FLC-coated exosomes to bind to antigenic determinants, we have compared the binding of wild type C57BL/6 mouse Ts cell exosomes, that are originally coated with Ab FLC (Figure 1C), to native OVA, OVA tryptic peptides and OVA 323-339 antigenic determinant, in an ELISA-based assay (Protocol, Figure 7F, left). This showed that the highest amount of the suppressive exosomes has bound to OVA tryptic peptides, then to native OVA and to OVA 323-339 peptide (Figure 7F). Taking into account that only a part of the OVA-primed B1a cells produce OVA-323-peptide-specific FLC, this observation unequivocally proved that the suppressive exosomes bind to antigenic determinant due to the coating with specific FLC, thereby allowing antigen-specific targeting of APCs expressing Ag peptide complexed with MHC for suppression of DTH response.

## 3. Discussion

### 3.1. Ag-specific Ts cell-derived exosomes mediate tolerance induced by Ag high dose administration

The current study has contributed greatly to further understanding of our original discovery of Ag-specific immunosuppressive exosomes [4]. These prior findings came from the hapten-specific effector T cell-mediated, in vivo system of cutaneous CS, where the Ag-specific tolerance was shown to be due to exosome-like nanovesicles coated with Ab FLC delivering miRNA-150 [4,8]. The central finding of this unique system was that systemic treatment of mice with such Ag-specific, immune suppressive exosome-like nanovesicles strongly inhibited established CS in actively sensitized hosts, by systemically administering single physiological doses of the vesicles at the 24-hour peak of the cutaneous immune response. Those inhibitory exosome-like nanovesicles were secreted by CD8^+^ Ts cells not expressing the Treg cell marker FoxP3 [4], and targeted Ag-presenting macrophages [6]. After induction of tolerance to hapten in donor mice, their suppressive exosome-like nanovesicles were found in their sera, plasma and in the supernatant of their cultured spleen and lymph node cells [4]. Thus, suppressive exosomes were released and acted systemically to achieve Ag-specific immunological tolerance, and were able to act at local sites of CS elicitation in the ear skin after systemic administration.

This current new study was extended to high dose OVA protein Ag-induced tolerance, that was similarly mediated by Ag-specific nanovesicles, determined to be common classical exosomes coated with anti-OVA Ab FLC, and also again delivering inhibitory miRNA-150, acting on macrophage APC, likely by binding to their surface expressed OVA Ag peptides in MHC to inhibit companion DTH effector T cells.

### 3.2. Development of the OVA protein Ag-induced DTH system regulated by Ag-specific, Ts cell-derived inhibitory exosomes delivering miRNA-150

Our findings in the hapten-induced CS system were new, unique, unconventional and a bit controversial. The findings were made in a system of hapten chemical Ag-specificity, and indicated that the exosome-like nanovesicles targeted the APC, guiding the eventual suppression of their companion CS-effector T cells [6]. However, the exact entity likely on the APC surface bound by the Ag-specific FLC coating these suppressive exosomes could not be determined in this CS system. This is because hapten immunization and challenge results in hapten binding over the whole APC surface as it becomes chemically covalently linked to diverse host proteins.

This current study was performed in the different, protein Ag-induced DTH system, and allowed expansion of the previous findings by being able to determine the Ag entity targeted on the APC. This enabled exosome-mediated transfer of miRNA-150, that in turn activated suppressive function of APC, that then inhibited companion OVA-Ag-specific DTH-effector T cells. Our new current findings indicate that peptides of the OVA Ag, likely in complex with MHC on the APC surface, are very likely bound by Ab FLC coating the surface of these Ts cell-derived suppressive exosomes.

There was a further important property of treatment with the previously studied hapten Ag-specific exosomes, expanded here in the OVA protein Ag DTH system. In CS, in vivo suppression was mediated by physiological doses of the nanovesicles, that was effective even when a single systemic IP exosome dose was administered at the in vivo maximum peak of the cutaneous response and lasted for several days [4]. In the current study, suppression of OVA DTH was achieved by systemic administration of a physiological dose of exosomes from OVA Ag-tolerized mice by the IP, IV, ID and even oral route; the latter producing the most powerful inhibition (Figure 3F).

### 3.3. Significance of administrating the suppressive exosomes orally

A remarkable new finding was that the OVA-specific suppressive exosomes can even be given orally to actively immunized recipients in a single dose, again at the height of the T cell effector response to achieve strong inhibition. In fact, this produced the strongest effect of all routes examined with prolonged suppression over four days of established DTH at the particular systemic site of the ear skin (Figure 3F). These findings are very important since in general they resemble the clinical situation of treating a patient with a pre-existing and ongoing immune-mediated reaction. Furthermore, successful and powerful suppressive treatment with exosomes via the oral route, if translated to the clinical situation, is a low impact and high effect treatment modality that obviously would be convenient to patients; especially children.

### 3.4. Hapten and protein Ag-specific suppressive exosomes

Our prior study was the first description of natural, Ag-specific exosome-like nanovesicles, uniquely composed of components from two different cell types; i.e. T cells and B cells. The T cell-derived component was miRNA-150 contained in exosome-like nanovesicles from hapten immune tolerance-mediating CD3^+^CD8^+^ Ts cells induced by administration of high systemic doses of hapten Ag [4]. These CD8^+^ Ts cell-derived exosomes had Ag non-specific, suppressive activity due to their ability to deliver inhibitory miRNA-150 to targeted cells [4]. Furthermore, anti-hapten B1a cells, also activated by the tolerogenic procedure, provided hapten-specific Ab FLC for coating the Ts cell-derived exosome-like nanovesicles to render them Ag-specific [4,8].

In the current study, OVA Ag-induced tolerance was also found to be mediated by CD3^+^CD8^+^ cells producing Ag-specific, suppressive exosomes, similarly coated with Ab FLC and particularly transferring miRNA-150 cargo. Further, the OVA-specific suppressive exosomes similarly targeted the APC companions of the finally suppressed OVA-specific DTH effector T cells (Figure 2B).

Other EV-mediated mechanisms of immune suppression have already been described and claimed to have Ag-specific activity. However, this has often been tested only in one-way Ag-activity system by simple comparison to challenge with another Ag or just media, or testing non-immune hosts with the involved Ag, as controls. Thus, the mechanisms in these studies should instead be considered only Ag-dependent [17-28]. Furthermore, the Ag-specificity of the function of other suppressive exosomes was not established in the important and meticulous study of RNAs involved in the suppression of Th1 cells by classical FoxP3^+^ Treg cells [25], and in another study on immunopathogenesis mediated by Treg cell-derived EVs [29].

In great contrast, our prior studies on CS suppression definitively established the strict hapten Ag-specificity of exosomes by dual reciprocal testing [4,8,10]. In the current study, we have also reciprocally tested the Ag-specificity of exosome-mediated tolerance to a pair of unrelated protein Ag; i.e. OVA and KLH. Accordingly, OVA-specific suppressive exosomes strongly inhibited effector DTH responses to OVA protein Ag in actively immunized mice. Reciprocally, KLH-specific suppressive exosomes failed to suppress OVA-induced DTH (Figures 3D and 3E). This proved that protein-Ag-specific suppressive exosomes acted in a strictly Ag-specific manner.

### 3.5. Protein Ag-specific, B1a cell-derived, non-suppressive exosomes can be rendered suppressive by in vitro association with miRNA-150

A very important aspect of the hapten CS-inhibiting, Ts-cell-derived exosomes was carriage and then delivery of miRNA-150 to targeted cells, that alone compared to controls inhibited Ag presentation by these acceptor macrophages [6]. Importantly, we confirmed this exact scenario in protein Ag high dose tolerance induced by exosomes mediating the suppression of OVA DTH, that was effective only in the presence of Ag-presenting macrophages. In both the CS and DTH circumstances, Ag-specific targeting of exosomes derived from CD3^+^CD8^+^ Ts cells was mediated by their surface Ab FLC derived from cells other than those that produced the exosomes (i.e. by co-activated B1a cells) [8].

Furthermore, we described another complex biological process that was based on the recognition that the essential aspects of the suppressive exosomes in CS and DTH was exosome surface Ag-specificity and delivery of miRNA-150 [4,10]. To construct this with natural components, we employed exosomes derived from B1a cells generated at only two days after ID immunization alone with OVA [11,30]. Although these B1a cell-derived exosomes were OVA Ag-specific due to the surface expression of OVA Ab FLC and/or anti-OVA BCR, they were not suppressive, since they lacked a miRNA-150 cargo, because the donors were not Ag-tolerized. Thus, of crucial significance, as in CS suppression [10], we found that in vitro association of these B1a cell-derived, OVA Ag-specific exosomes with miRNA-150 alone rendered them suppressive of the OVA-specific effector T cells via the APC. Further, these miRNA-150-associated, anti-OVA B1a cell-derived exosomes were strongly suppressive over subsequent days, when injected systemically at the 24 hour peak of the OVA-specific DTH response (Figure 5F), and this was comparable to suppression mediated by the OVA-specific Ts cell-derived exosomes (Figures 3D and 3F) [10]. In summary, we hold that achieving this suppression by simply combining the naturally produced B1a cell-derived exosomes from optimally immunized donors, that express surface Ab, together with selection of carried miRNA-150, is the strongest proof of principal definitively verifying the ideas concerning the biologically functional suppressive exosomes. This became the main interest of our ongoing studies.

### 3.6. Dose response testing to determine the limiting dose of associating miRNA-150 to mediate suppression of OVA DTH by B1a cell exosomes

This in vitro procedure of associating miRNA-150 with anti-OVA B1a cell-derived exosomes allowed as well for testing the limiting dose of associating miRNA-150 able to mediate suppression of OVA DTH. Such a rarely done dose response experiment showed that the minute dose of only 300ng or 1×10^−13^ moles of miRNA-150 sufficed for suppression of an OVA DTH effector cell mixture adoptively transferring DTH (Figure 5E). This compares to the very unusual, ultra-low dose of 50×10^−15^ moles, or 50 femtomoles of miRNA-150, associating with B1a cell-derived hapten Ag-specific exosomes for suppression of CS adoptive cell transfer [10]. However, there is as yet no method allowing accurate determination of the percentage of miRNA-150 taken up by these exosomes and further exosome uptake by the targeted cells. Therefore in actuality, much less miRNA-150 may be exosome-transferred in both cases to induce the biological effects.

The apparent difference in limiting dose of miRNA-150 in CS and DTH may be due to the distinctive aspects of the involved mechanisms. In CS the targeted cells are exposed to and linked with many molecules of reactive hapten applied to these macrophage APC, and thus likely have great amounts of hapten covalently conjugated to multiple entities on their surface beyond conjugating to self peptides complexed in MHC. This would allow for great diversity of Ab-linked suppressive exosome binding, uptake, and possible subsequent induction of intracellular mechanisms mediated by transferred miRNA-150. In contrast, in OVA DTH, similar Ab-coated suppressive exosomes just seem to bind the foreign Ag OVA-derived peptides only expressed in complexes with limited numbers of MHC molecules on the APC surface, as the only possible Ag-specific targets for their surface Ab FLC.

### 3.7. Confirmation that Ag-specific exosomes mediate miRNA delivered suppression in two different DTH systems

Various findings of Ab-dependent, exosome-mediated suppression in hapten CS, were now repeated in protein Ag DTH. As a first potential treatment scenario, this offered a unique situation of constructing functionally suppressive exosomes derived again from two cell types; i.e. T cells and B cells. Firstly, this has potential clinical significance, since there is the tolerogenic route of inducing Ts cell-derived endogenous miRNA-containing exosomes to be coated by either native or chosen B1a cell-derived, endogenously available Ab FLC.

In a second scenario, aimed at creating suppressive B cell-derived exosomes for treatment, immunization of patients with exogenous Ag or altered self-Ag might induce a B1a cell source of exosomes that are Ag-specific due to either a surface expression of BCR specific for that immunizing Ag, or via a coating of endogenous Ab FLC. Additionally, two exosome types; i.e. Ts cell- or B1 cell-derived, that could be, respectively, endogenously or in vitro associated with a chosen miRNA, or likely siRNA, would then mediate selected miRNA-dependent alteration of APC function. Indeed, when transferred to totally syngeneic recipients (Figures 3F and 5E), they may induce desired epigenetic alterations in specifically targeted host APC, that bear peptide surface determinants expressing the given Ag to be bound by the Ab FLC or Ab-like BCR on the exosome surface. The in vitro miRNA association process would follow steps in the new alternate pathway for in vivo exosome-mediated transfer of extracellular RNA (exRNA) between cells, that we described previously [10]. Clinically, there are other possibly applicable conditions, that involve protein Ag DTH-like systems; possibly including protein immune reactivity in: asthma, autoimmunity, transplantation or cancer. Since exosomes seem to be universal nanoparticles of all of life, these constructs would constitute a new simple approach for simultaneous and mutually interacting Ag and gene-specific therapy to curb excessive DTH or related T cell reactivity in such disease processes.

### 3.8. Potential application of dual Ag- and gene-specific exosome to clinical conditions like anti-cancer therapies

Similar combined specific treatment of leukemia and cancers is becoming available in emerging chimeric T cell antigen receptor (CAR) therapy. This might be delivered instead by exosomes with advantages as cell-free carriers, a far less complicated modality for targeted therapy in cancers and possibly in other diseases [31]. However, CAR therapy is dependent on a single V-region gene encoded T cell reactivity to a single given peptide-Ag/MHC determinant. Unfortunately, this fixed property is subjected to the well-known ability of cancers to evolve resistance to a given therapy, in this case away from the vulnerable single tumor expressed monoclonal T cell receptor for Ag-peptide/MHC [32-34]. This compares to the possibility to use exosomes coated with easily rotated Ab of differing specificity and also to the dual possibility to use readily selected, different therapeutic miRNA or siRNA. Together this achieves varying Ag or oncogene specific exosome-mediated therapy, likely much less prone to tumor escape.

Furthermore, in a variety of cancers, treatment with anti-PD-1 or anti-PD-L1 Ab, or other analogous negative co-stimulation specific Ab, that often is incomplete when used as Ab alone, could be used as an exosome coating for check point targeting to reverse tumor mediated suppression of host effector T cell anti-tumor reactivity. When armed with multiple check point-targeting mAb, such therapeutic, Ag-specific exosomes might not only cover multiple target check points, making tumor escape less likely, but also could simultaneously deliver relevant inhibitory RNAs to disable the function of involved suppressive cells [35]. Accordingly, exosomes could be constructed to inhibit cancer cell-induced immunosuppression by both, specific anti-PD1/L1 Ab coating and delivery of engineered hybridizing gene sequences encoded by selected DNA or RNA, for instance in CRISPR/cas-9 oncogene targeting [36]. Additionally, for these various scenarios of dual Ag and gene specific exosome-mediated anti-cancer therapies, the prospect of efficacious oral therapy would undoubtedly have greater patient acceptance and comfort, especially when combined with existing drug and radiotherapy that might be reduced to achieve less toxicity in the patients.

### 3.9. Ab FLC surface binding is associated with “activated” exosomes

A unique feature of both, the hapten-Ag and OVA protein-Ag systems, is that the generated exosomes are able to bind selected Ag-specific Ab FLC and associate with chosen miRNA-150. These unusual binding and associating features uniquely provide both highly Ag-specific binding to targeted cells and ability to transfer gene directed, functional alterations; such as epigenetic changes. Such dual-specific targeting, in part resulting from cargo transfer, is a unique property of the exosomes we have described, as no other laboratories have yet reported these combined properties. In contrast, when exosomes are similarly prepared from normal, unmanipulated animals, they do not have either significant Ab FLC coating or miRNA-150 associating properties compared to those from animals stimulated by diverse modes of positive immunization or immune tolerization.

We refer to these exosomes having the combined Ab FLC binding and miRNA associating properties as “activated” exosomes, as they are preferentially derived from lymphoid cells of immunized or tolerized mice, and not unmanipulated normal donors [4,10]. It is currently not known what steps in immunization or tolerization induce these activation properties, nor what are the stimuli that produce these changes, that are not found on exosomes from non-immune donors. Acquisition of activation characteristics is true for exosomes of mice undergoing strong and prolonged exposure to high doses of Ag in tolerogenesis, or even mere two day ID immunization with proteins without an adjuvant, and to exosomes from various Ag high dose-tolerized Ab-deficient or miRNA KO mice as well.

The association of the mAb FLC was determined by ability to bind the surface of “activated”, but not normal, exosomes. This was verified by cytometric visualization of FLC on the exosome surface (Figures 1C and 5B) [4,8]. Such surface and other activations seemed to account for acquisition of both, the ability to bind specific Ag via surface Ab FLC, that enabled mediation of Ag-specific suppressive function, and also to associate and then transfer selected miRNA for alteration of intracellular functional effects.

This Ag-specificity was demonstrated by OVA Ag-affinity column separation of the total suppressive exosome population that yielded two subpopulations (Figure 2A) [4]. These were specifically Ag-binding, Ab-coated, therefore considered “activated”, classical small exosomes that had the suppressive functional activity vs. Ag-non-binding and functionally inactive exosomes (Figure 2A). This suggested that, in this case, when employing Ag-affinity column separation, only around 10% of the total exosomes from immunized or tolerized animals is actually “activated” and binds the Ag [4].

Prior studies with mast cells originally determined binding of Ab FLC to the cell membrane, that, as shown here, also applies to the surface of “activated” exosomes. The binding of Ab FLC to the mast cell surface enabled subsequent binding of specific Ag [37]. This allowed for induction of mast cell degranulation and mediator release [37,38], and correlated with the activity of mast cells in several models of immunological diseases [39]. Another connected and also unusual property of the “activated” exosomes was that there was no bioactive exosome surface binding of Ab free heavy chains, nor whole Ag-specific IgG of the same Ab source, despite their ability to bind the Ab FLC [4]. This was also true for the mast cell binding [37,38].

### 3.10. Possible role of membrane lipids in EV surface activation

We have formed the hypothesis that exosome activation is due to an increased content of components in their membranes. This idea is based on our unpublished findings that these vesicles have greater size (150-225 nm) than the typical exosome-like nanovesicles (50-120 nm), and by flow cytometry showing that they can have lower expression of routine molecules present on the surface of most exosomes, like MHC class I (unpublished). This suggests that there has been a filling in of an expanded surface with membrane components, that we believe are likely lipids. This would be the most immediately manipulatable surface component during exosome donor cell stimulation [40]. Further, study of cellular uptake of exosomes showed that interactions with targeted cells can be modified by changing the lipid composition of their membranes [41,42].

Concerning membrane lipids and Ab FLC binding, our ideas also came from studies showing that binding of immunoglobulin FLC to human mononuclear cell subsets depends on surface lipids [43-45]. These ideas were sustained by our preliminary unpublished studies examining lipid binding of FLC. Firstly, mAb FLC were shown to have stronger binding ability to certain individual lipid components, such a phosphoethanolamine. This was determined firstly with an ELISA-type assay of lipids adsorbed on plastic wells and then bound by mAb FLC, quantitated by a second binding with anti-light chain mAb linked with enzyme and detected enzymatically. Secondly, liposomes constructed at small EV size and dominantly composed of the best individual lipid binders, like phosphoethanolamine, and less of poor Ab FLC binders like sphingomyelin, were able to bind the Ab FLC, as detected by Western blotting. Thirdly, these Ab FLC-associated liposomes had biological properties that routinely composed liposomes did not have (Hutchinson and Askenase, unpublished data).

Studies are underway to similarly examine miRNA interactions with activated vs. normal exosomes. The amounts of miRNA involved in association with “activated” OVA suppressive exosomes are likely minute, as we showed before in CS [10], and here in DTH (Figure 5E). It was hoped that biotinylation of miRNA-150 would be helpful for such analysis, since it could aid detection of the binding of miRNA to the exosome surface or associated within the “activated” exosomes by pull down techniques [46]. Unfortunately, this biotinylation caused inactivation of miRNA-150 biological properties (Figure 5E, Group G); as predicted in the literature [47].

### 3.11. Similar “activated” exosomes may mediate RNA transfers in other systems

Lipid changes of “activated” exosome membranes may account for their other exceptional properties, like our unique finding that oral systemic suppressive therapy with Ag-specific exosomes can affect a strong local immune reaction at a specific cutaneous anatomical site. This may relate to the physiologically transferred functions of numerous exosomes particularly contained in mothers’ milk and delivered to neonates [48-50]. The special property of these exosomes is resistance to harsh conditions in the neonatal stomach, like very low pH plus digestive enzymes [51,52], that also could be properties of “activated” exosomes in the immunological systems we described [53]. RNA cargo of milk exosomes may transfer epigenetic information to neonates after intestinal absorption [54,55], perhaps via intestinal epithelial cell endocytosis [56,57], benefitting the neonatal intestine [58,59], and allowing postulated subsequent systemic transfer [60], that may influence neonatal immunity [48,49,61-64], bone formation [65,66], and development of the nervous and endocrine systems in the neonates. However, there are as yet few studies actually showing beneficial systemic effects in neonates of orally-delivered milk EV-carried miRNAs, and there is some contrary data [67]. Thus, our findings of systemic oral exosome transfer of miRNA mediated immune suppression strengthens the idea that milk exosomes might orally transfer systemic effects to the neonates.

Of further great relevance, there are numerous claims that among miRNAs taken orally in foods, in some cases these mixtures contain exosomes that also can pass stomach digestion for uptake systemically to affect host processes [68-70]. Again, there is controversy with some compelling data against this concept [67,71-74]. Note, however, that these are complex issues with many technical barriers to be overcome for definitive determinations before this can be settled [75-78]. Herein however, oral treatment with claimed “activated” subsets of exosomes carrying miRNA-150 in our non-dietary system clearly showed strong and prolonged in vivo immune suppression at a particular systemic skin site. Thus, suppressive effects of orally administered, Ts cell-derived, Ag-specific exosomes (Figure 3F), delivering a particular inhibitory miRNA-150, favor the idea of transferred functionality of dietary food miRNAs, under some circumstances.

### 3.12. Demonstration that Ag-derived peptides are the binding targets of FLC on Ag-specific suppressive exosomes

Our prior study in the CS hapten-system showed that macrophage APC and not the finally suppressed effector T cells were the direct target of the Ag-specific suppressive exosomes [6]. We hypothesized that Ab FLC on the surface of the inhibitory exosomes bound to Ag peptides in MHC complexes on the APC surface to lead to eventual suppression by the miRNA-150-influenced APC of the effector T cells that mediate CS [30].

Our success in generation of OVA protein Ag-specific suppressive exosomes made testing of these ideas possible. Firstly, we showed binding of Ts cell-derived exosomes to OVA-coupled sepharose on an Ag-affinity column only by effectively suppressive exosomes (Figure 2A), and to an anti-CD9-coupled Ab-affinity column (Figure 1F), that strongly supports our hypothesis. In addition, cytometric analysis revealed the co-expression of CD9 and Ab FLC by exosomes from both Ts and B1a cells (Figures 1C and 5B). To further extend these findings, we set up an ELISA with native OVA absorbed on plastic wells, and determined the binding of four different anti-OVA mAb rendered into FLC, as well as the isolated heavy chains, compared to whole anti-OVA IgGs. We found that binding of OVA by IgG was the strongest, heavy chains next and FLC last, but definitely detectable (Figures 6A and 6B). Comparing the four mAb, there was a consistent hierarchy of binding for all chains and whole IgG among all. Thus, binding native OVA by each mAb and its chains was different; ranging from excellent to much less, and to nil or nearly nil (Figure 6). The similar binding within a mAb IgG and its heavy and light chains likely was due to identical immunoglobulin V-region DNA sequence mutations in the producing B cells, and thus identical specificity of the translated mAb. Conversely, the different binding between the four mAb likely was due to different V-region mutations resulting in different derived mAb affinities for native OVA Ag determinants adsorbed on plastic.

Knowing that Ag-presenting macrophages were the target of the Ab FLC on the exosomes for mediating the final Ts cell suppression of the effector T cells [4,6], we further tested suppression by the four mAb FLC separately coating the same exosomes from Ag-tolerized, Ab-deficient JH^neg/neg^ mice, that gave a remarkable result. We observed a very different hierarchy in the strength of mAb dependent exosome-mediated suppression (Figure 7B) compared to binding of native OVA (Figures 6A and 6B). This was consistent with the idea that the Ab FLC on the suppressive exosomes bound to Ag-peptides on the APC and not to native OVA.

To investigate this further, we made a tryptic digested peptide mixture from native OVA and tested its ability to block suppression mediated by these exosomes, using different protocols. In the first experiments, the OVA peptides were incubated in vitro with the exosomes coated with two different anti-OVA mAb FLC that gave intermediate strength suppression. In both cases we found strong reversal of suppression, when FLC-coated exosomes were blocked by OVA tryptic peptides (Figure 7E). Additionally, from past studies we knew that the two day sera of OVA-immunized mice contain both anti-OVA IgM and also Ab FLC, derived from activated B1a cells [9,16,79]. Therefore, we also coated JH^neg/neg^ mouse Ts cell-derived exosomes with this sera, presumably containing polyclonal immune anti-OVA Ab FLC (Figure 6E), that in fact produced strong suppression (Figure 7D). Inhibitory blocking of suppression by these polyclonal Ab FLC with OVA tryptic peptides resulted in the best reversal of suppression (Figure 7E, Groups G and H); likely due to the polyclonal nature of the serum Ab vs. the mAb FLC. This overall result contributed significant evidence suggesting that Ab FLC coating the Ag-specific exosomes bound to OVA Ag peptides on the APC surface. This would confer the Ag-specificity of Ts cell-mediated immune tolerance [8]. Noteworthy, we have recently reported that, in the particular circumstances, self-tolerant exosomes can be redirected toward suppression of allergen-induced DTH responses by coating with allergen-specific Ab FLC [80].

In a second protocol, the OVA peptide mixture was used to construct peptide Ag-affinity columns. Passage of the Ab FLC down this column enabled testing those that actually had bound peptide on the column in the Ag-specific exosome-mediated suppression of DTH. The same two mAb FLC, intermediately active when coating exosomes for Ag-specific suppression, and serum polyclonal anti-OVA Ab FLC, were applied to OVA peptide columns. The OVA peptide binding Ab FLC fractions were eluted from each column and then used to coat exosomes from Ag tolerized JH^neg/neg^ mice in the suppression assay. These eluted FLC, that had bound the peptide mixture, when subsequently coating these exosomes, led to strong suppression of DTH (Figure 7D). This result confirmed that those Ab FLC that can bind peptides on the column, may mediate the Ag-specificity of the suppressive exosomes (Figure 7D, Groups C, D and E) [8].

This unanticipated result can be seen as further evidence that the binding of the anti-OVA Ab FLC on the suppressive exosomes is likely to OVA peptides complexed in MHC on the APC surface. Thus, an explanation may be due to the manner of binding peptides by Ab FLC in the ELISA compared to FLC on the exosome surface that bind peptides in MHC. In the ELISA, there is only binding of low-affinity, isolated Ab FLC dispersed in solution to the peptides adsorbed on the artificial plastic surfaces in non-physiological conformations. In comparison, the in vivo measured, Ag-specific exosome-mediated suppression depends on FLC that are not free but linked to the exosome surface. Here, the mAb FLC are not dispersed but in multiples closely together on the exosome surface, turning a series of low affinity bindings into far greater avidity for binding Ag determinant of physiological conformation when complexed with MHC, which was confirmed in vitro in ELISA-based assay (Figure 7F). However, more direct study of mAb FLC binding to the peptide-MHC complex will be needed to prove this point in vivo.

### 3.13. Summary of the new findings of biological and clinical significance of the Ag- and miRNA-specific exosomes that suppress OVA protein-induced DTH

New findings brought by our current studies could be summarized into several main points. (i) Uniquely, suppressive function depends on exosomes consisting of components from two lymphocyte types (T and B cells). (ii) Ag-specificity in DTH suppression is due to exosome surface coating with anti-OVA Ab FLC. (iii) This Ag-specificity was definitively demonstrated by reciprocal functional suppression testing of two distinct protein-specific suppressive exosomes. (iv) Also, there was definitive demonstration that T or B cell exosome-mediated suppression in DTH is mediated by delivered inhibitory miRNA-150. (v) Systemic suppression of DTH could be achieved in vivo by administration of single physiological doses of exosomes; remarkably including oral administration. (vi) Strong, systemic suppression of maximal in vivo DTH responses at a given peripheral tissue site was observed over several days after administration of a single physiological dose of Ab FLC coated and miRNa-150 delivering exosomes. (vii) Exosomes suppressing DTH could be constructed with chosen dual Ag-specific targeting and the transfer of selected miRNA-encoded epigenetic functional effects by mixing and matching chosen surface Ab FLC and selected miRNA. (viii) These exosomes mediated suppression by very low miRNA-150 associating doses. (ix) Determination that the Ab FLC coating the suppressive exosomes likely binds OVA peptides that are complexed in MHC on the APC surface was shown firstly by comparing in vitro binding of the Ab FLC to OVA, compared to strength of suppression when coating exosomes from Ag-tolerized, Ab-deficient JH^neg/neg^ mice; and secondly by different peptide inhibition protocols.

To summarize, Ag-specific, Ab FLC-dependent, “activated” exosome delivery of suppressive miRNA-150 via targeted binding of Ag-peptides in MHC on APC provides a new mechanism of immune regulation with great clinical significance, since it employs natural physiologic exosomes that can be given orally and fashioned for both chosen Ag-specific targeting and selected miRNA-specific functional alterations of targeted cells that can last for several days.

## 4. Materials and Methods

### 4.1. Mice

Nine to ten-week-old CBA/J, BALB/c, JH knock out (JH^neg/neg^), C57BL/6 or miRNA-150 knock out (miRNA-150^neg/neg^) male mice from Jackson Laboratories (Bar Harbor, ME) or CBA mice from the Breeding Unit of the Jagiellonian University Medical College, Faculty of Medicine (Krakow, Poland) were fed autoclaved food and water ad libitum. All experiments were performed according to the guidelines of Ethics Committees of Yale (approval No 07381) and Jagiellonian (approval No 122/2013 and 88/2017) Universities.

### 4.2. Antigens, reagents and culture media

The following reagents were used: ovalbumin (OVA), serum-free Mishell-Dutton medium, RPMI-1640, minimal essential medium with amino acids, HEPES, 2-mercaptoethanol (Sigma-Aldrich, St Louis, MO), keyhole limpet hemocyanin (KLH, Stellar Biotechnologies), Dulbecco’s phosphate-buffered saline (DPBS), penicillin/streptomycin, sodium pyruvate, L-glutamine (Gibco Life Technologies, Grand Island, NY), acetone, ethanol, glucose (P.O.Ch., Gliwice, Poland), EDC (1-ethyl-3-(3-dimethylaminopropyl)-carbodiamide, Pierce, Thermo Fisher Scientific, Waltham, MA), EDTA (BDH, Poole, UK), extra virgin olive oil (Basso Fedelee Figli, San Michele di Serino, Italy).

### 4.3. Generation of Ts cell-derived, protein Ag-specific, suppressive exosomes

Ts cells from OVA Ag-tolerized mice releasing suppressive exosomes were induced in an analogous manner to the tolerogenic procedures described in CS [4]. Thus, mice were injected IV on days 0 and 4 with 0.2 ml of a freshly prepared 10% DPBS suspension of syngeneic erythrocytes conjugated with OVA or, in some instances, with KLH protein Ag. We determined in preliminary experiments that EDC is an efficient activating agent for the coupling of protein Ag to erythrocyte membrane proteins. For full tolerogenesis these IV treatments were followed by intradermal (ID) immunization on two consecutive days 8 and 9 with a total of 0.2 ml of a 0.5 mg/ml OVA each time (100 µg, injected into four sites in the abdominal areas at 25 µg per site), or similarly KLH, both used as a 0.9% NaCl-solution and without adjuvant. After lymph node and spleen collection on day 11, single cell suspensions were cultured at 37°C in protein-free Mishell-Dutton medium at a concentration of 2×10^7^ cells/ml for 48 hours. To enrich for exosomes, the resulting culture supernatant was subsequently centrifuged at 300g and then 3,000g for 10 min, then filtered through 0.45- and then 0.22-micrometer molecular filters (Miltenyi Biotec) and finally ultracentrifuged twice at 100,000g for 70 minutes at 4°C [4]. The resulting pellet was resuspended in DPBS and used as OVA Ag-specific (or KLH Ag-specific) exosomes [4]. This pellet containing OVA-specific Ts cell-derived exosomes was absorbed onto copper grids and negatively stained with 3% uranyl acetate, and then visualized under transmission electron microscope (JEOL JEM2100, Tokyo, Japan). Furthermore, it was subjected to Nanoparticle Tracking Analysis (NTA, Nanosight) [4].

In some instances prior to culture, mouse lymph node and spleen cells from Ag-tolerized donors, were depleted of either CD3^+^ or CD8^+^ cells by incubation with, respectively, anti-CD3 or anti-CD8 monoclonal IgG antibodies and rabbit complement for 60 minutes in 37°C water-bath. Afterwards, dead cells were removed by discontinuous gradient centrifugation on Ficoll (1.077g/mL, GE Healthcare).

In some experiments, actively tolerized mice were ear challenged on day 11, and subsequent ear swelling responses were measured as described below. In some instances, tolerized and actively immunized mice were depleted of clodronate-sensitive cells (i.e., macrophages) by IP injection of 0.2 ml of clodronate liposomes (Department of Molecular Cell Biology, Vrije University, Amsterdam, The Netherlands) in PBS, at 24 hours before ear challenge [6].

### 4.4. In vivo treatment of mice actively immunized with OVA with suppressive exosomes injected at the peak of DTH response

Mice were immunized ID with a total of 100 µg of OVA, as described above, and 5 days later DTH ear swelling was elicited by ID injection of 10 µl of OVA 0.9% NaCl solution (0.5 mg/ml, thus 5 µg per ear) into both ears. The pellet containing OVA Ag- or KLH Ag-specific suppressive exosomes was used for treatment of the mice actively ID immunized for elicitation of OVA-specific DTH either at immunization, or just before ear challenge or at the peak of the response 24 hours after ear challenge with OVA Ag, at a dose of 1×10^10^ EVs per mouse [11]. These mice were systemically IP injected with the DPBS suspension of exosomes at a dose of about 1×10^10^ nanovesicles per recipient. This model allowed testing of Ag-specificity of exosome action. Subsequent ear thickness was measured with an engineer’s micrometer daily up to 120 hours after challenge by a blinded observer [4,12].

To calculate the increase in ear thickness indicating strength of DTH, values of background ear thickness, measured before ear challenge, were subtracted from values of ear thickness at particular time points after ear challenge. Furthermore, to evaluate the net swelling response, mean ear thickness increase in non-immunized but similarly OVA Ag challenged littermate control mice was subtracted from the ear thickness increase measured in each ear of mouse of each control or experimental group. In general, groups consisted of 5 mice and experiments were repeated 2-4 times. The average ear swelling was expressed as the delta ± standard error (SE), after subtraction of the negative control value. The two-tailed Student t test or one-way Analysis of Variance (ANOVA) with post hoc RIR Tukey test were used to assess the significance of differences between groups, with p values of less than 0.05 taken as the minimum level of statistical significance.

Exosome counts were estimated by Nanoparticle Tracking Analysis (Nanosite). The dose used for treatment of actively immunized mice was considered physiological, since this is the average number of exosomes per ml of blood in normal unmanipulated mice.

Where indicated, exosomes either derived from cultures of Ts-cells or derived from early two day OVA immune B1a cell-derived supernatants and then associated in vitro with inhibitory miRNA-150, were systemically administered, at the same doses, but in different volumes of DPBS (0.2-0.5 mL), directly to actively immunized mice via IV, IP or ID routes. Further and most importantly, the same suppressive exosomes at the same dose were also administered orally (per os, PO) using a gastric feeding tube. For all of the compared systemic exosome treatments, suppressive and control exosomes were administered just after measurement of the 24 hour peak of active ear swelling response. In the case of PO delivery of the exosomes, mice were kept fasting for 2 hours before and 1 hour after administration of the suppressive exosomes.

### 4.5. In vitro treatment of OVA DTH effector cell mixture with exosomes, prior to their adoptive transfer to naive recipients

OVA DTH-effector cells were obtained from mixed spleen and lymph node cells of mice actively sensitized by ID multiple injections of a total of 100 µg OVA in plain 0.9% NaCl (as described above) and harvested at day 4 after immunization [4,11]. OVA Ag-specific Ts-cell-derived, suppressive exosomes were used at the same dose of 1×10^10^ EVs for in vitro pulsing of a mixture of DTH-effector T cells and APC, at 7×10^7^ total lymphoid cells to be transferred per eventual recipient. Then, the exosome pulsed DTH effector cells were adoptively transferred into naive recipients, in which 24 hours later DTH ear swelling responses were elicited by ID injection of 10 µl of OVA 0.9% NaCl solution (0.5 mg/ml, thus 5 µg per ear). Ears were measured for thickness with an engineer’s micrometer (Mitutoyo, Japan), at 24 hours after challenge. Results were statistically analyzed as described above.

### 4.6. Polyribonucleotide treatments to induce suppressive activity of non-suppressive exosomes, or to block their suppressive activity, prior to incubation with DTH effector cell mixture

In some instances, Ts-cell-derived, OVA-specific exosomes were pretreated with a polynucleotide antagonist of miRNA-150 (anti-miRNA-150, Dharmacon, Lafayette, CO), at a dose of 3 µg per total exosomes (10^10^) per each of eventual five recipients of exosome-pulsed DTH-effector cells (7×10^7^, the average number of cells transferred IV to each recipient). After washing away of unbound anti-miR-150 molecules by ultracentrifugation, the mixture of DTH effector cells and miRNA antagonist associated exosomes were incubated in a 37°C water-bath for 30 min. Then, the exosome treated and control DTH effector cells were adoptively transferred into naive recipients, as described above.

In other experiments, exosomes from Ag-tolerized miRNA-150^neg/neg^ mice were treated in a 37°C water-bath for 30 minutes with synthetic miRNA-150 in a dose of 3 µg per 10^10^ exosomes to be used subsequently to treat the DTH-effector cell mixture (7×10^7^), per single eventual recipient. Then, through ultracentrifugation, we removed excessive, exosome-non-associated miRNA-150 molecules and used the resulting miRNA-150 pulsed exosomes to treat the DTH effector cells in vitro prior to washing away the exosomes and then IV adoptive transfer of the cells.

### 4.7. Ab FLC coating, OVA peptide blocking, and OVA Ag or anti-CD9 affinity column chromatography of Ts-cell-derived exosomes

Ts cell-derived exosomes from Ag tolerized JH^neg/neg^ mice were non-suppressive due to the lack of surface FLC, but nonetheless carried miRNA-150. These were converted for Ag-specific suppression by surface coating with various anti-OVA FLC (derived from either four different monoclonal Ab or immune serum of 2 day OVA-immunized mice) by incubation overnight on ice. This was followed by ultracentrifugation at 100,000g to remove unbound FLC into the supernatant from the Ab FLC-coated exosomes in the pellet.

In other instances, OVA-specific Ts cell-derived exosomes endogenously coated with B1a cell-derived Ab FLC, were initially blocked with tryptic digested OVA peptides by a preliminary 2 hour incubation in 37°C water-bath. This was followed by ultracentrifugation to remove excessive OVA peptides into the supernatant from peptide-blocked exosomes in the pellet. In yet other cases, these OVA-specific, Ts-cell-derived exosomes were separated by affinity chromatography on columns filled with sepharose coupled with either native OVA Ag or with purified anti-CD9 monoclonal antibodies [4].

### 4.8. Generation of B1a cell-derived exosomes and their in vitro association with miRNA-150

To induce very early activated Ag-specific B1a cells, mice were immunized with OVA protein Ag by ID injections of total volume of 0.2 ml for 4 separate simultaneous intradermal injections of a 0.5 mg/ml OVA solution in 0.9% NaCl (total ID immunizing dose of 100 µg OVA) without adjuvants on days 0 and 1 [11]. Then, on day 3, lymph nodes and spleens containing the immune B1a cells were collected and single cell suspensions were cultured in protein-free Mishell-Dutton medium at a concentration of 2×10^7^ cells/ml for 48 hours [9]. The resulting culture supernatant was subsequently centrifuged at 300g and then 3,000g for 10 min, filtered through 0.45- and then 0.22-micrometer molecular filters and then ultracentrifuged twice at 100,000g for 70 minutes at 4°C. The pellet presumably containing B1a cell-derived exosomes was absorbed onto copper grids and negatively stained with osmium tetroxide, and then visualized under transmission electron microscope (FEI Tecnai T12, LaB6).

The resulting pellet containing enriched Ag-specific B1a cell-derived exosomes was resuspended in DPBS. Then, where indicated, the Ag-specific B1a cell-derived, non-suppressive exosomes that received no endogenous miRNA-150, since they were from immunized and not tolerized mice, were associated in vitro with synthetic miRNA-150. This was achieved by incubation in 37°C water-bath for 30 minutes with miRNA-150 at a dose of 3µg per 10^10^ nanovesicles. These exogenously miRNA-150-associated exosomes were subsequently used to pulse 7×10^7^ DTH-effector cells per single recipient. Prior to incubation with the DTH-effector cells, the Ag-specific, B1a cell-derived exosomes that had been incubated with free miRNA-150 for association, were washed by a single ultracentrifugation to remove excessive, exosome-non-associated miRNA-150 molecules.

### 4.9. Cytometric analysis of exosomes and evaluating the presence of antibody light chains in 2 day serum of OVA-immunized mice

Aldehyde/sulphate latex beads (4 µm, Life Technologies, Thermo-Fisher Scientific, Carlsbad, CA) were incubated in DPBS with OVA-specific exosomes derived from Ts cells or B1a cells at room temperature for 2 hours with gentle agitation. Afterwards, exosome-coated beads were blocked with 100 mM glycine, washed, resuspended in DPBS, and stained with fluorescein isothiocyanate (FITC)-conjugated mAb against mouse kappa light chains (BD Biosciences, San Diego, CA) and/or phycoerythrin (PE)-conjugated mAb against mouse CD9, CD63 or CD81 tetraspanins (BD Biosciences, San Diego, CA). Alternatively, the aldehyde/sulphate latex beads were coated with native OVA Ag by overnight incubation at 4°C with gentle agitation, and then, after washing, were incubated with 2 day immune serum from the ID OVA-immunized mice. After washing, these OVA-coupled beads were stained against mouse antibody kappa light chains to evaluate the eventual presence of anti-OVA FLC in the serum.

### 4.10. ELISA for assessment of binding to native OVA, OVA tryptic peptides or OVA 323-339 peptide by various anti-OVA mAb FLC

Isolated OVA-specific mAb FLC and isolated mAb free heavy chains were made from four different OVA-specific mAb IgGs (clones F2 3.58, 7E9.17, 7H7.E1, and 11B.1) from the Frank Fitch collection, University of Chicago, that were kindly provided by Anne Sperling. The whole IgGs were reduced and alkylated to separate heavy and light chains in the laboratory of Frank Redegeld at the University of Utrecht, The Netherlands. Proteins were eluted from columns with 6M guanidine and subsequently loaded onto a gel filtration column (HiLoad 16/60 Superdex 200pg, GE Healthcare) to separate anti-OVA mAb heavy and light chains (FLC). Subsequently, samples were dialyzed versus PBS to remove guanidine and then concentrated.

Then, OVA Ag-specific mAb FLC, heavy chains, and whole IgGs were added to quadruplicate microwells of high binding ELISA plates that previously were coated with native OVA (25 µg/mL). After Ag coating, the plates were blocked with PBS/10% FBS/0.05% Tween-20 for 1 hour at room temperature, washed and finally incubated with the four separate OVA-Ag-specific Ab FLC, heavy chains, or whole IgG1, all diluted in PBS/1%FBS/0.05% Tween-20 (assay buffer) at multiple serial dilutions for 1 hour at room temperature. For detection of bound mAb FLC, heavy chains and whole IgGs, plates were subsequently incubated with 0.1 µg/mL HRP-labeled goat anti-mouse κ light chain Ab (Southern Biotech, Birmingham, AL 35260), diluted in assay buffer for 1 hour at room temperature. For detection of mAb heavy chains, 0.1 µg/mL HRP-labeled goat anti-mouse IgG/IgM H+L Ab (Jackson Labs) was used. Finally, TMB was used as a substrate for the enzymatic reaction, terminated by adding 1 M H_2_SO_4_ after 23 minutes in the case of Ab FLC assaying or after 3 minutes in all other cases. Between incubation steps, wells were washed three times with PBS/0.05% Tween-20. To evaluate the binding of FLC to OVA 323-339 peptide, plate wells were coated with OVA 323-339 peptide at a concentration of 10 µg/mL, and assay was developed as above. To compare binding of FLC to native OVA and OVA tryptic peptides, plate wells were coated with OVA preparations at a concentration of 100 µg/mL, and the assay was performed as above, but the reaction was stopped by adding 1 M H_3_PO_4_ after 8 minutes.

### 4.11. ELISA for assessment of binding to native OVA, OVA tryptic peptides or OVA 323-339 peptide by either JH^neg/neg^ mouse exosomes coated with anti-OVA mAb FLC or wild type mouse exosomes

Plate wells were coated with either native OVA, OVA tryptic peptides or OVA 323-339 peptide, all at a concentration of 100 µg/mL overnight at 4°C and then blocked with 2% BSA for 2 hours at room temperature. Then exosome samples (approximately 5×10^9^ exosomes per well) were added to particular wells and incubated overnight at 4°C. After washing with 0.1% BSA in PBS, 100 µL/well of rat monoclonal IgG antibody anti-mouse CD9 (BD Biosciences) was added at a concentration of 4 µg/mL and plate was incubated for 2 hours at room temperature. After washing, 100 µL/well of HRP-conjugated rabbit anti-rat IgG antibody (Invitrogen, ThermoFisher) was added at a concentration of 2 µg/mL and plate was incubated for 1 hour at room temperature. After extensive washing, 100 µL/well of TMB substrate was added and the reaction was terminated after 8 minutes by adding 50 µL/well of 1 M H3PO_4_.

## Supplementary Materials

Supplementary materials can be found at www.mdpi.com/xxx/s1.

## Author Contributions

Conceptualization, K.N., K.B., W.P. and P.W.A.; methodology, K.N. and K.B.; validation, K.N., K.B. and P.W.A.; formal analysis, K.N. and K.B.; investigation, K.N., K.B., T.G.K.; data curation, K.N. and K.B.; writing—original draft preparation, P.W.A.; writing—review and editing, K.N. and K.B.; visualization, K.N., K.B., and P.W.A; supervision, K.B. and P.W.A.; project administration, K.B. and P.W.A.; funding acquisition, K.B. and P.W.A. All authors have read and agreed to the published version of the manuscript.

## Acknowledgments

We are indebted to Dr. Peter Cresswell (Yale) for his advice and guidance in the APC and peptide aspects of this work, to Dr. Joan Steitz (Yale) of her advice on the miRNA aspects, and to Dr. Bernadeta Nowak (Jagiellonian University Medical College) for her valuable advice on flow cytometry analysis of exosomes. We are very grateful to Dr. Rafal Szatanek (Jagiellonian University Medical College) for supporting Nanoparticle Tracking Analysis of the vesicles as well as to Dr. Olga Woznicka (Jagiellonian University), and to Lab Members of Electron Microscopy Unit (Yale School of Medicine) for their precious help with transmission electron microscopy of the vesicles. Results showed in the Figures 1A, 1B, 2A, 3A, 3B, 3D, 3F, 4B, 5D, 5E, 5F, 6E and 7F were obtained from experiments supported by Polish National Science Centre (NCN, 2013/11/B/NZ6/02041).

## Abbreviations

Ab: Antibody
Ag: Antigen
APC: Antigen-presenting cell
CAR: Chimeric antigen receptor
CS: Contact sensitivity
DTH: Delayed-type hypersensitivity
ELISA: Enzyme-linked immunosorbent assay
exRNA: Extracellular ribonucleic acid
EVs: Extracellular vesicles
FLC: Free light chains
ID: Intradermal
IP: Intraperitoneal
IV: Intravenous
KLH: Keyhole limpet hemocyanin
OVA: Ovalbumin
OVA-RBC: Ovalbumin-coupled red blood cells
OX: Oxazolone
PO: Per os, oral administration
RBC: Red blood cells
TNP: Trinitrophenol
Treg cell: Regulatory T cell
Ts cell: Suppressor T cell
WT: Wild type

## Supplementary figures S1 and S2

**Figure S1.**
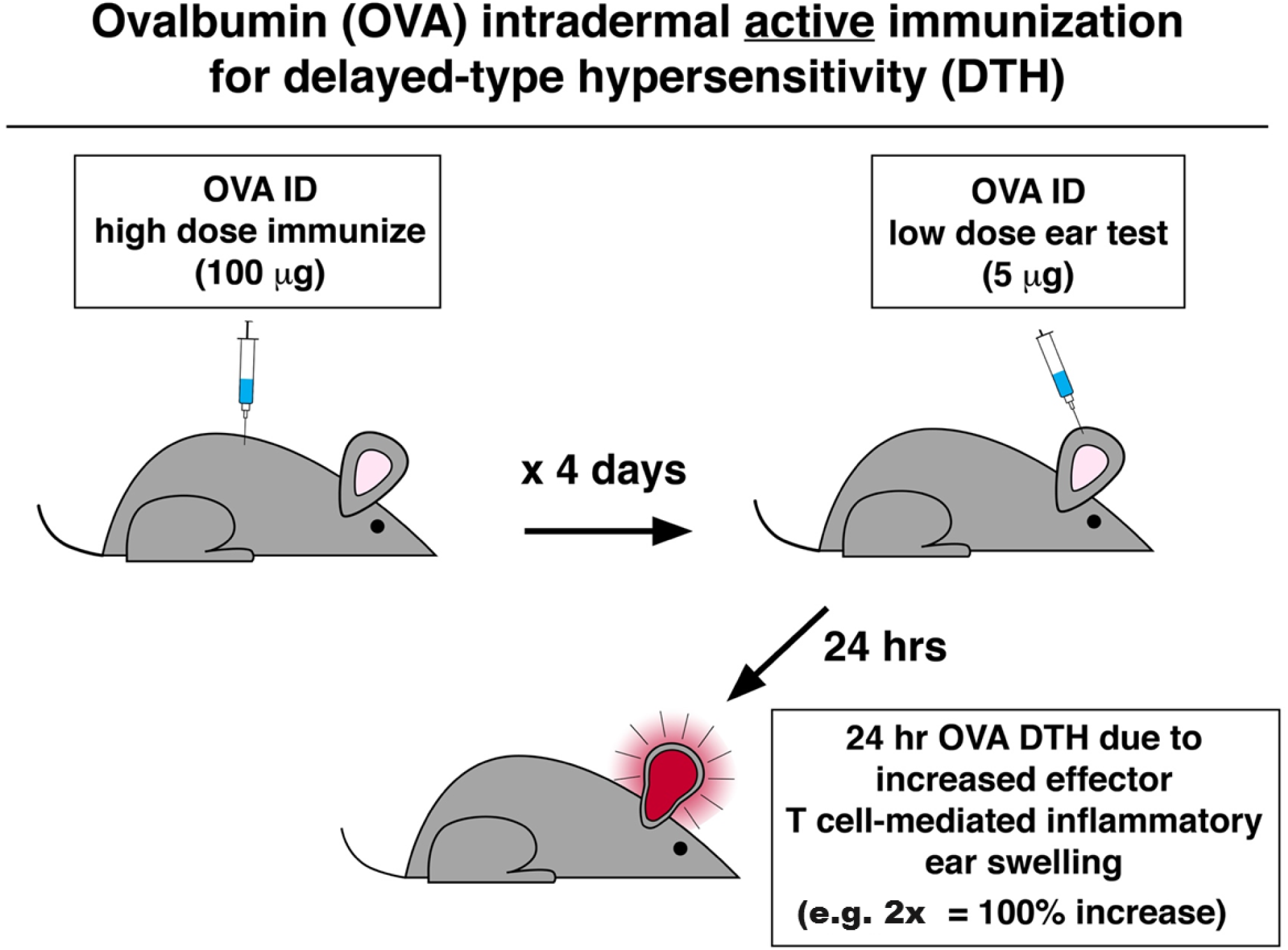
The general scheme of Ag immunization and challenge testing of mice administered ID with ovalbumin (OVA) without an adjuvant to induce DTH responses. **(S1A)** <u>Protocol for ID immunization and eliciting ear swelling responses after inducing OVA Ag DTH</u>. CBA male mice were immunized ID twice with OVA in saline injected into four different sites of the abdomen in 50µL each, for a total dose of 100µg OVA. Four days later they were skin challenged ID in the ears with 5µg OVA and the subsequent ear swelling was measured with an engineer’s micrometer at 2 and 24 hours, and, where indicated, daily up to 120 hours after challenge, by a well experienced reader, unaware of their experimental status. The data were recorded by an assistant who randomly selected mice for measurements and administered ether inhalation anesthesia for the ear readings. All readings were done in duplicate for each of both ears to obtain the single mean ear swelling determination for that mouse, at that single time point. Mice were kept fed and watered in a bacteriology hood in the laboratory for the duration of the experiment to protect them from infection and from the biological changes that might happen during transit to and from the animal quarters far away.

**(S1B).**
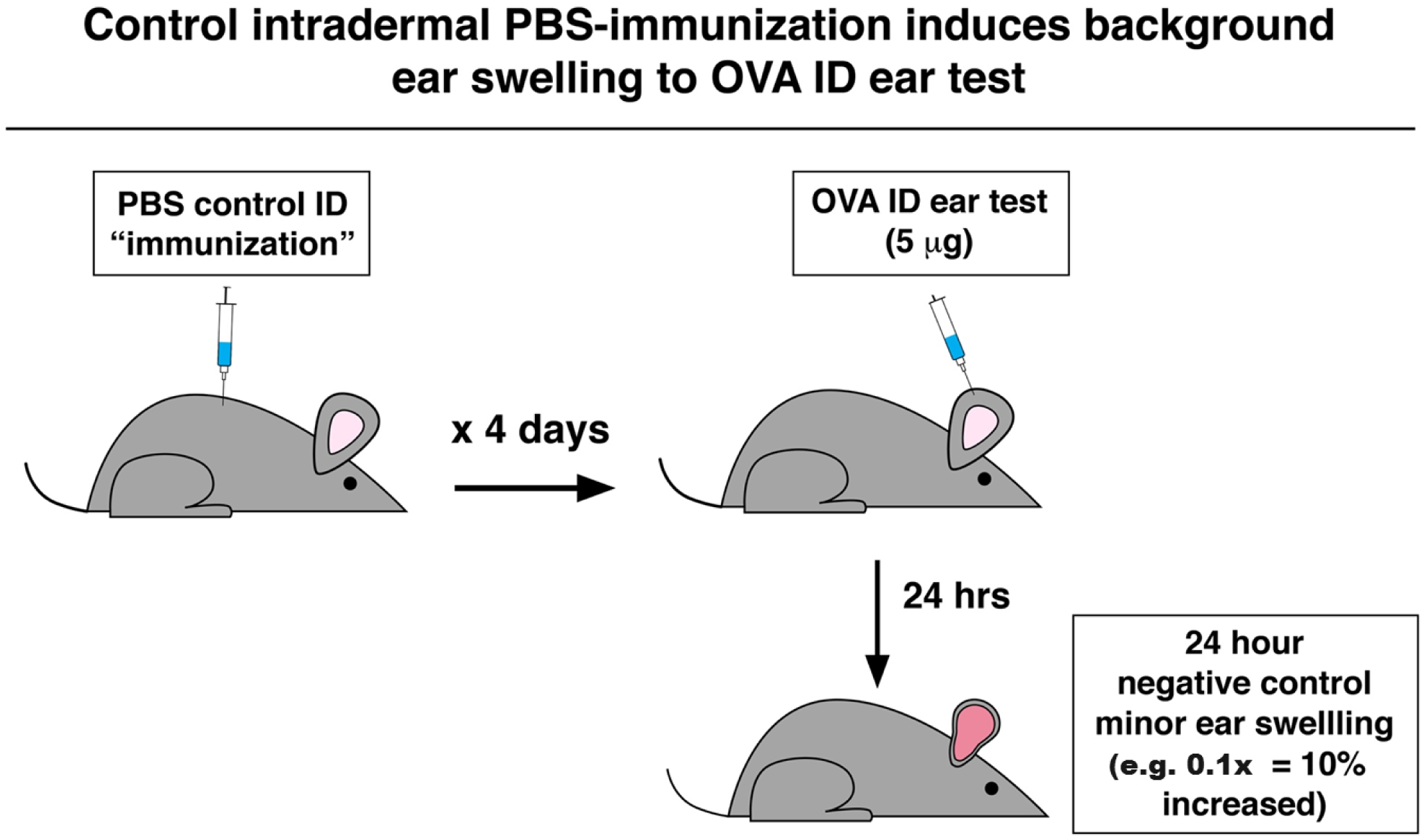
Protocol for control ID immunization and OVA skin testing to verify OVA DTH. Immunization is as in **Figure S1A**, but with ID saline alone. Then, four days later, ear challenge was exactly as above with 5µg OVA ID. Ear thickness measurements were exactly as above in **Figure S1A**.

**(S1C).**
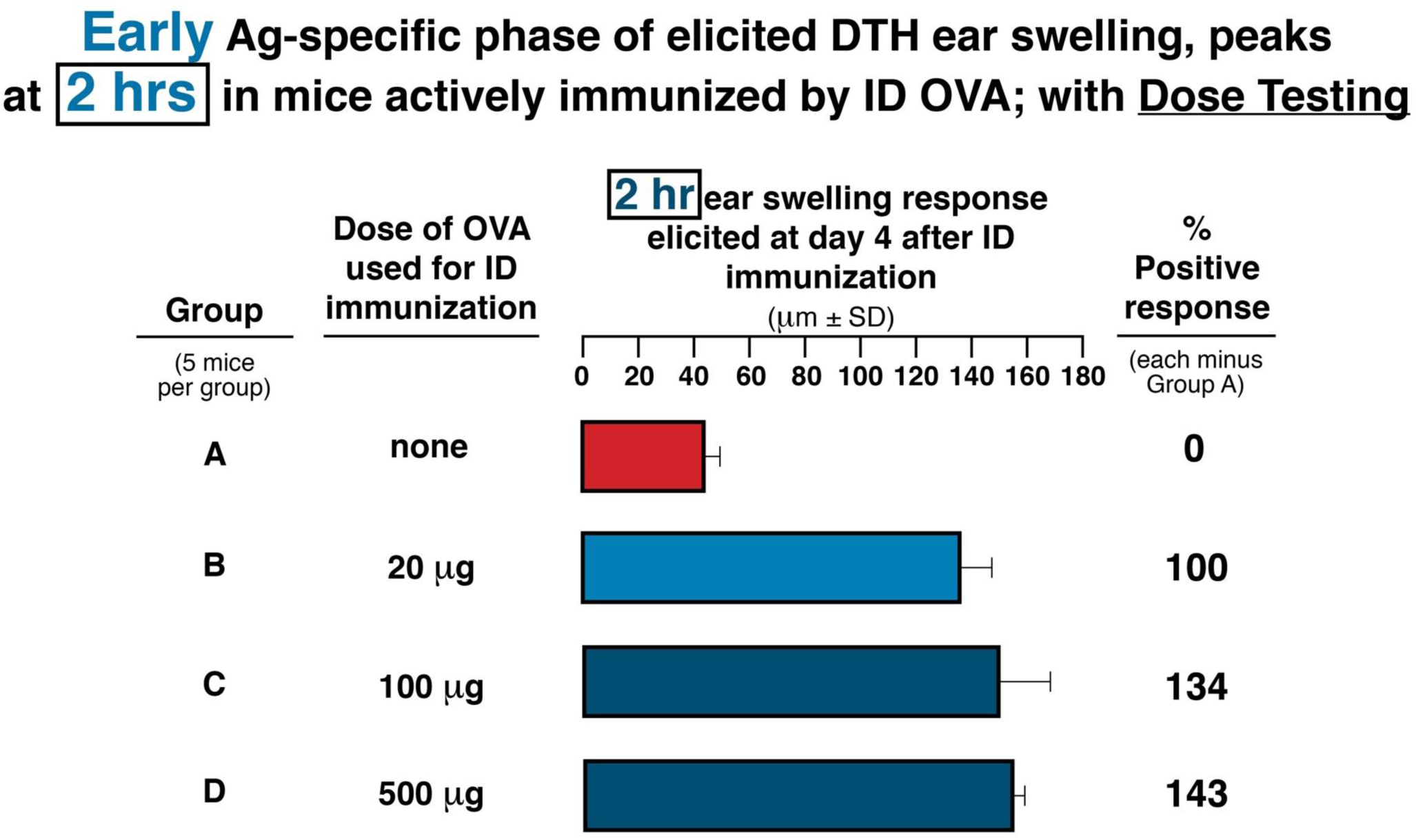
Weakly Ag-specific early phase of DTH is elicited as ear swelling that peaks at 2 hours after challenge with ID injection of OVA into the ears of mice actively immunized ID on day 0 and 1 with OVA; performed here with immunizing dose response testing. Immunization was as in **Figure S1A**, with ID total immunizing doses in each separate mouse were 20 µg, vs. 100 µg, vs. 500 µg OVA at four ID sites each. Then at day four, the 5 µg OVA ID elicited ear swelling responses were tested at 2 hours to evaluate an early phase of DTH.

**(S1D).**
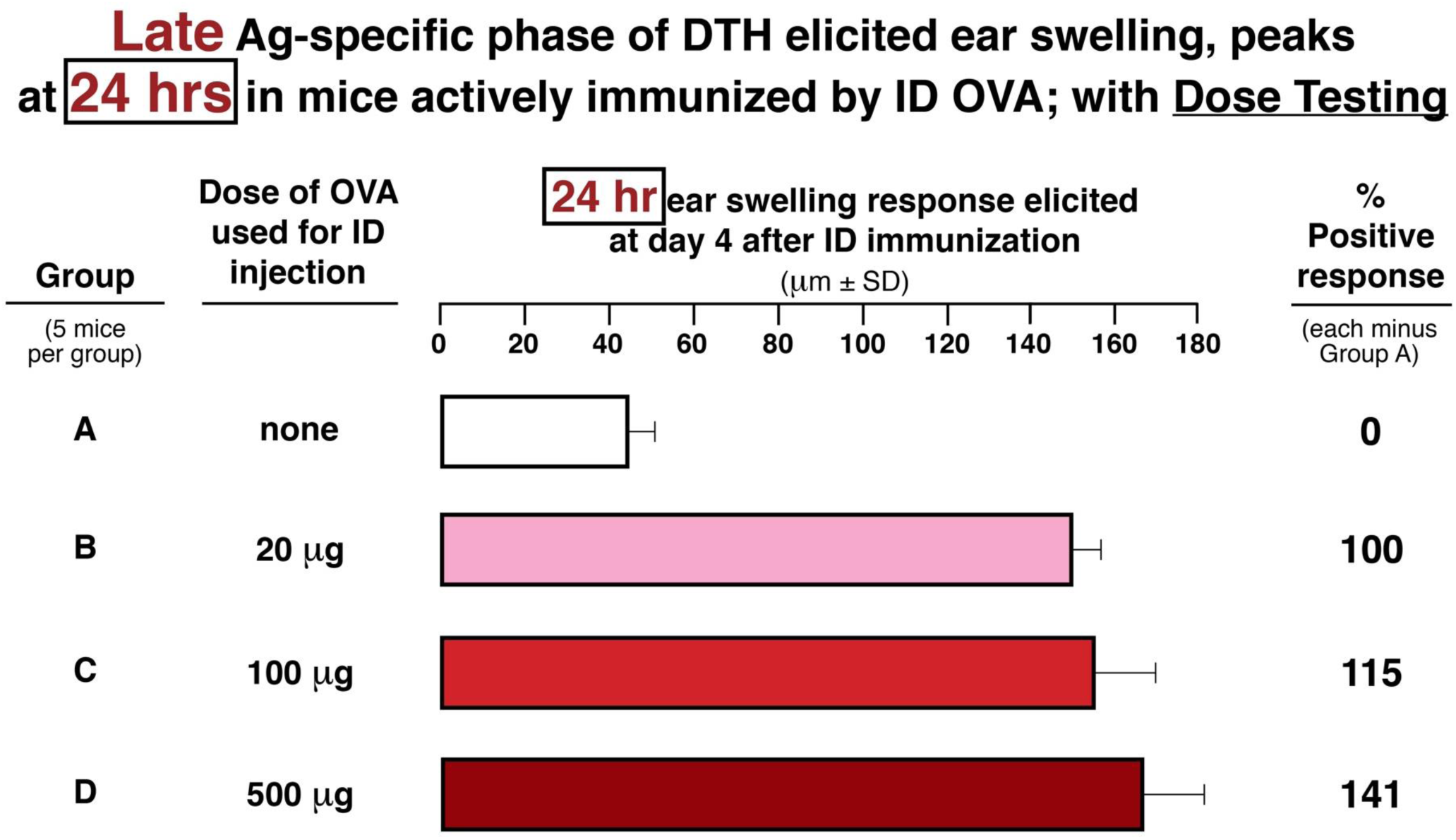
Late Ag-specific phase of DTH-elicited ear swelling response to local ID OVA injection, peaks at 24 hours in mice actively immunized ID with OVA; performed here with immunizing dose response testing. The ear swellings at 24 hour after challenge testing of mice from **Figure S1C**.

**Figure S2.**
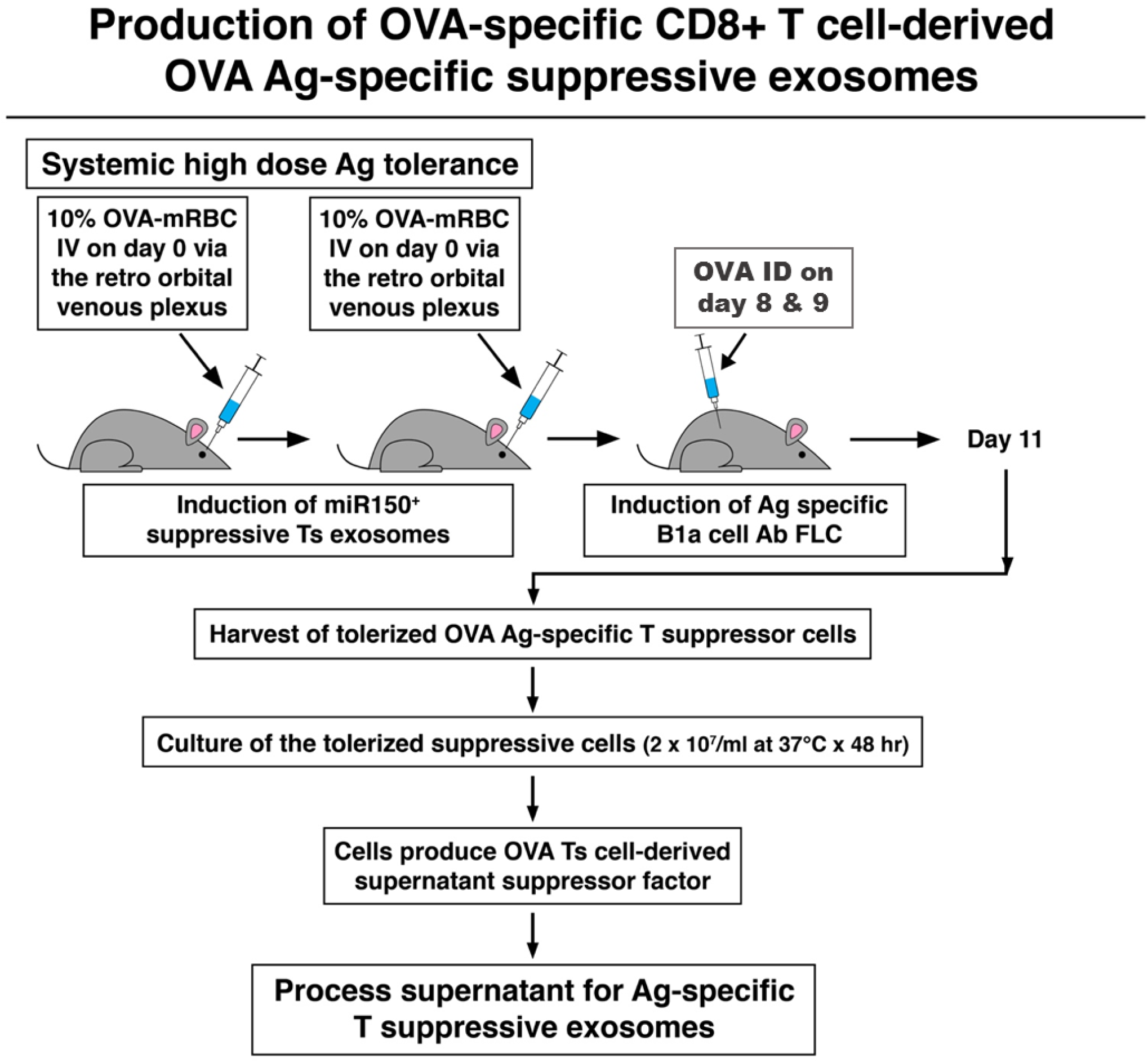
DTH induced by ID immunization with OVA is prevented by prior IV injection of high doses of OVA-linked autologous RBC that induce tolerance to OVA, due to suppressor T cells producing exosomes able to inhibit adoptive cell transfer of OVA-immune DTH. **(S2A)** <u>Protocol for induction of OVA Ag-specific suppressor T cells and their suppressive exosomes.</u> To induce immunological tolerance, mice were injected IV with a high dose of 10% autologous RBC-linked with OVA on day 0 and day 4. Then, on day 9, these mice received ID injections of OVA alone, as in **Figure S1A**. Then, on day 11, harvested lymph node and spleen cells were shown to contain suppressor T cells. The cells then were cultured in vitro at 37^°^C in protein free media at high density (2×10^7^/ml) for 48 hours and then the resulting suppressive supernatant was processed to obtain the contained exosomes, i.e. the supernatant was cleared of debris and cells by light centrifugations ending at 3,000g, then was ultrafiltered down to 0.2 µm (200 nano Meter) filters and subsequently 4°C ultracentrifuged twice at 100,000g to obtain a pellet rich in the exosomes (**Figure S1A)**.

**(S2B).**
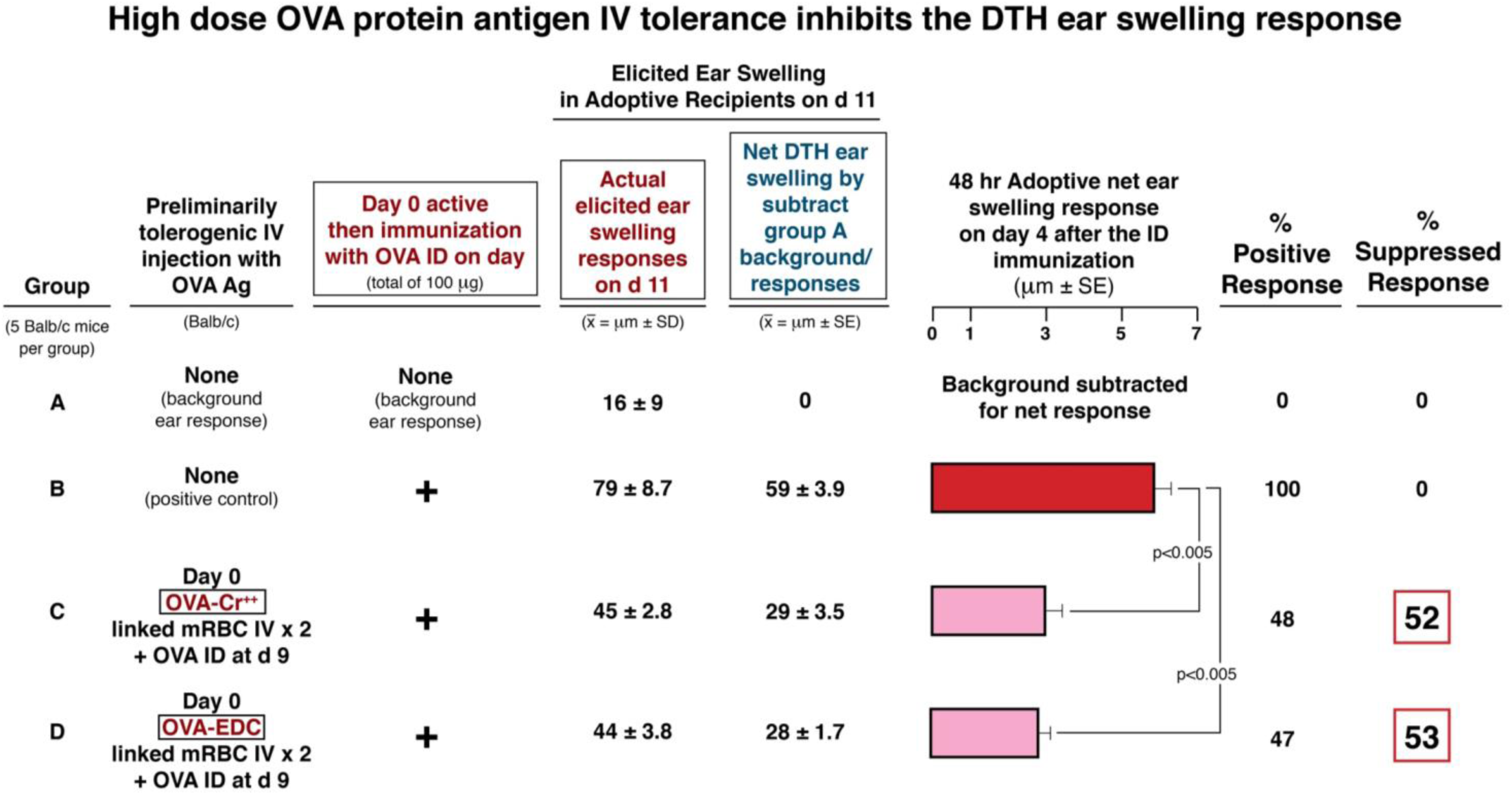
DTH ear swelling 4 days after ID immunization with OVA in mice pretreated with OVA-linked to autologous RBC by different reagents. Mice were injected IV twice with 10% autologous RBC as above, after linkage of the RBC to OVA by chromium or EDC (Groups C and D). Then on day 9 these mice, and previously non injected positive controls (Group B) were ID immunized with OVA alone as in **Figure S1A**. Four days later all groups, and an additional group of totally naive background mice (Group A) were ear tested as above, with 5 µg OVA ID and elicited DTH ear swelling responses were measured.

**(S2C).**
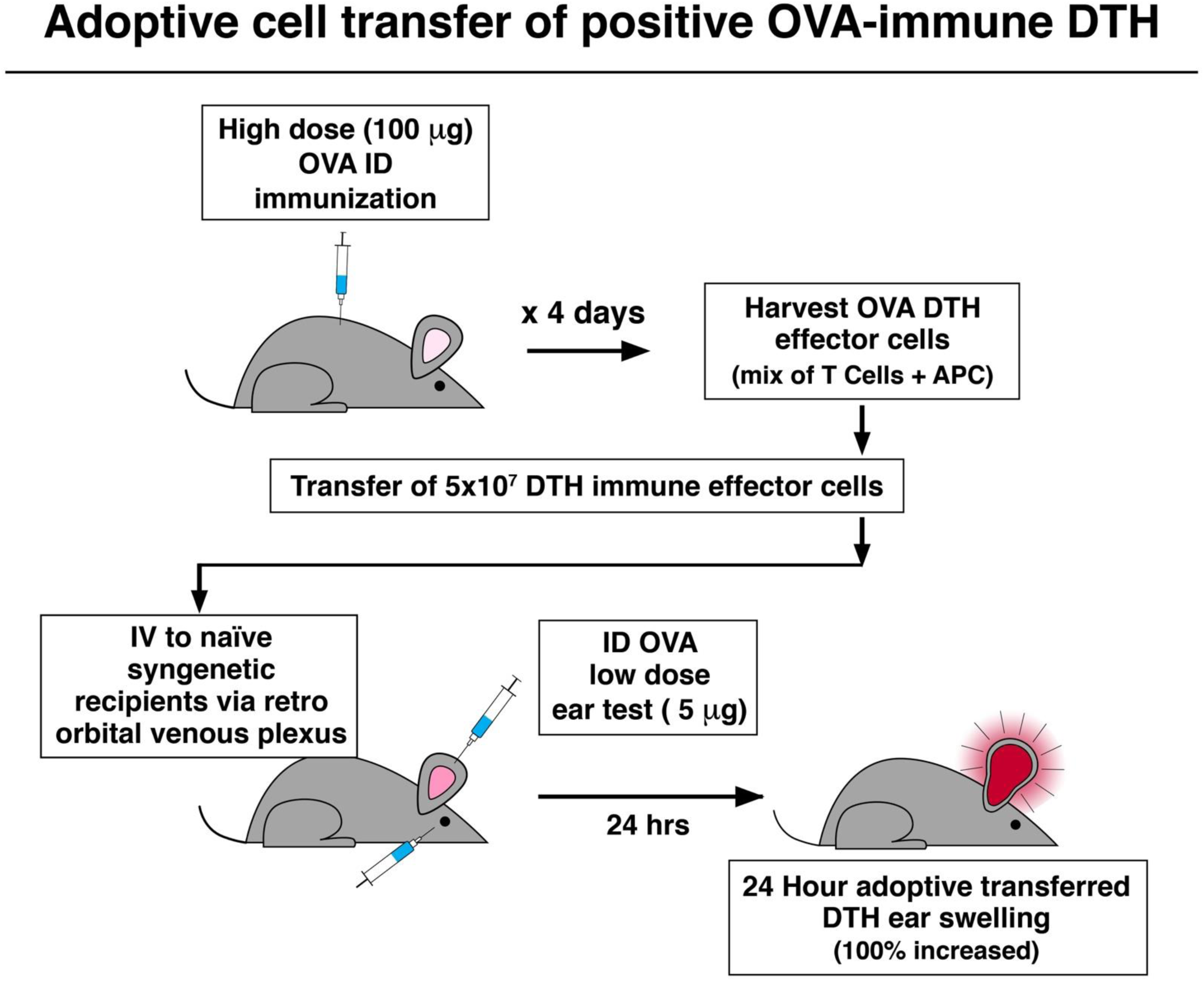
Protocol for adoptive cell transfer of OVA-immune effector cells from ID skin immunized donors to elicit DTH responses in recipients. Cell donor mice were ID immunized twice with 100 µg OVA in saline, as in **Figure S1A**. On day 4 lymph node and spleen cells containing DTH-effector cells were harvested. Then their single-cell suspensions were washed and subsequently resuspended in a volume of 0.25 ml/eventual recipient that was injected IV into naive mice. A day later the recipients were ear tested with OVA injected ID to elicit DTH ear swelling, that was measured in the recipients 24 hours later.

**(S2D).**
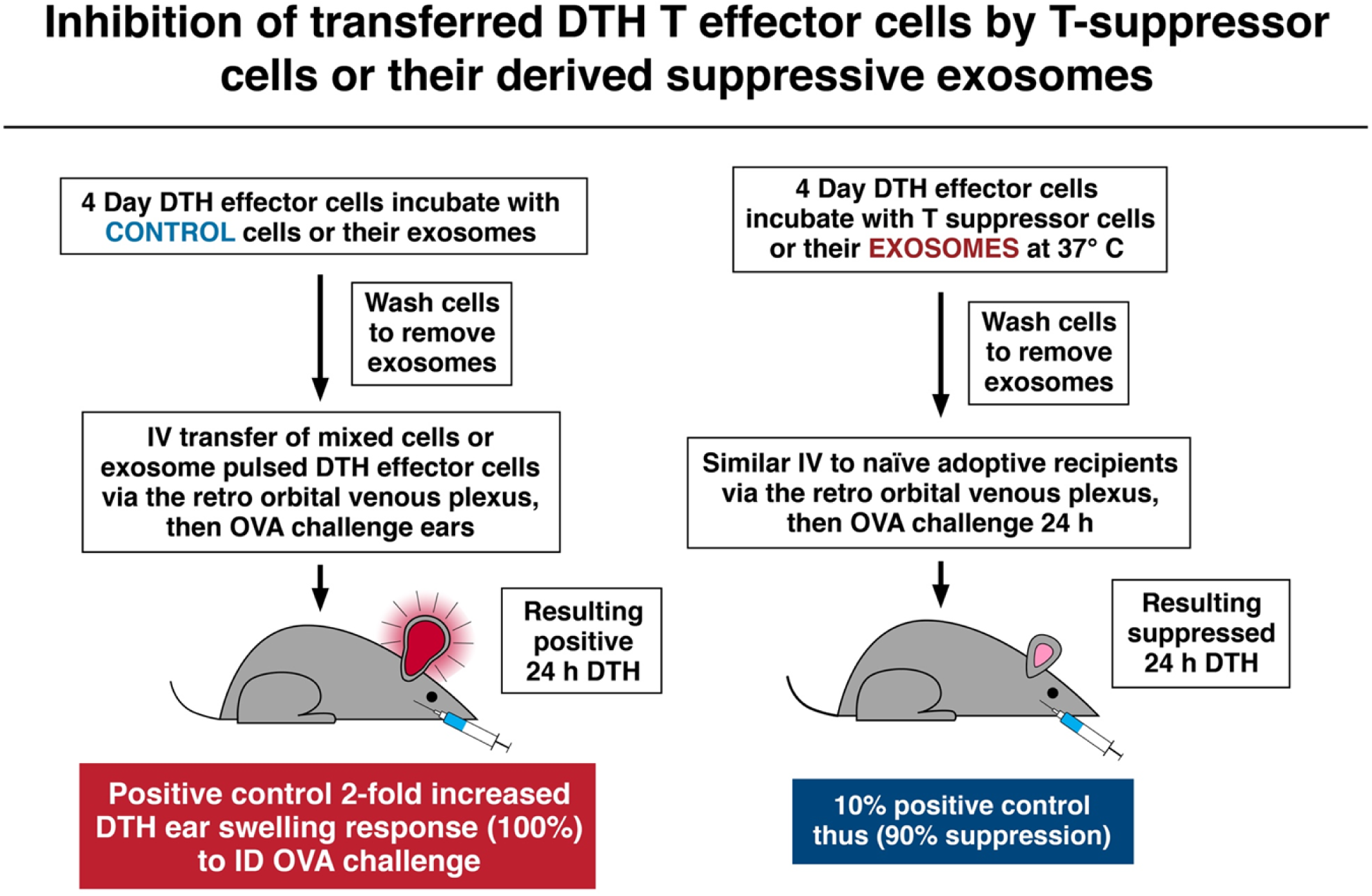
Protocol for inhibition of adoptively transferred DTH-effector cells by suppressor T cells derived from OVA-tolerized mice, or their derived suppressive exosomes. Adoptive transfer of DTH-effector cells was performed as above. However in these cases, the harvested effector cells were incubated in vitro at 37°C for 30 minutes with the suppressor T cell-derived exosomes, at a ratio of 10^10^ nanovesicles per 7×10^7^ effector cells to be transferred per eventual recipient. Then, after washing, the cells freed of the exosomes by centrifugation at 300g were transferred IV. The control transfers are shown on the left, and the experimental groups on the right of the figure.

